# Myco- and photobiont associations in crustose lichens in the McMurdo Dry Valleys (Antarctica) reveal high differentiation along an elevational gradient

**DOI:** 10.1101/718262

**Authors:** Monika Wagner, Arne C. Bathke, Craig Cary, Robert R. Junker, Wolfgang Trutschnig, Ulrike Ruprecht

**Author notes:** Corresponding Author: Ulrike Ruprecht, 0043-662-80445519.

## Abstract

The climate conditions of the McMurdo Dry Valleys (78° S) are characterized by low temperatures and low precipitation. The annual temperatures at the valley bottoms have a mean range from −30 °C to −15 °C and decrease with elevation. Precipitation occurs mostly in form of snow (3-50 mm a^−1^ water equivalent) and, liquid water is rare across much of the landscape for most of the year and represents the primary limitation to biological activity. Snow delivered off the polar plateau by drainage winds, dew and humidity provided by clouds and fog are important water sources for rock inhibiting crustose lichens. In addition, the combination of the extremely low humidity and drying caused by foehn winds, confined to lower areas of the valleys, with colder and moister air at higher altitudes creates a strongly improving water availability gradient with elevation.

We investigated the diversity and interaction specificity of myco-/photobiont associations of a total of 232 crustose lichen specimens, collected along an elevational gradient (171-959 m a.s.l.) within the McMurdo Dry Valleys with regard to the spatial distribution caused by climatological and geographical factors. For the identification of the mycobiont and photobiont species three markers each were amplified (nrITS, mtSSU, RPB1 and nrITS, psbJ-L, COX2, respectivley). Elevation, associated with a water availability gradient, turned out to be the key factor explaining most of the distribution patterns of the mycobionts. Pairwise comparisons showed *Lecidea cancriformis* and *Rhizoplaca macleanii* to be significantly more common at higher, and *Carbonea vorticosa* and *Lecidea polypycnidophora* at lower, elevations. Lichen photobionts were dominated by the globally distributed *Trebouxia* OTU, *Tr_*A02 which occurred at all habitats. Network specialization resulting from mycobiont-photobiont bipartite network structure varied with elevation and associated abiotic factors.

Along an elevational gradient, the spatial distribution, diversity and genetic variability of the lichen symbionts appear to be mainly influenced by improved water relations at higher altitudes.

## Introduction

The McMurdo Dry Valleys (MDV) in Southern Victoria Land of Continental Antarctica are characterized by an environment that is exceptional also for Antarctica: it is extremely arid and cold, which makes it hostile for most organisms. Thus, life is rare within the valleys of this polar desert, and only few life forms can cope with these extreme conditions (e.g. Adams et al. 2006; Pointing et al. 2009). The main limiting factor for life within the MDV is water availability with fog, clouds, dew and ephemeral melting water of snow patches having important effects on the climatic conditions (Adams et al. 2006; Green et al. 2007; Pannewitz et al. 2005; Stichbury et al. 2011). Among the most diverse macro-organisms present in the MDV are lichens. Lichens represent a classic example of symbiosis, consisting of a fungus (mycobiont) and one or more photosynthetic partners (photobiont). When completely desiccated, lichens are dormant and can survive unfavorable conditions for long periods (Green 2009; Kappen and Valladares 2007). As a consequence they are able to colonize rocks and boulders above melting streams or in the vicinity of snow patches, even in such extreme environments as the MDV (e.g. Green et al. 2011b; Hertel 2007; Ruprecht et al. 2012a; Ruprecht et al. 2010; Sancho et al. 2017; Schroeter et al. 2010). The most successful species are green-algal lichens, as they do not depend on the presence of liquid water for reactivation and can be active below zero degrees, in contrast to cyanobacterial lichens that appear to be completely absent in continental Antarctica (Green et al. 2011a; Kappen 2000; Lange et al. 1986; Schlensog et al. 1997; Seppelt et al. 2010).

Several studies on mycobiont-photobiont interactions in lichens have shown that green-algal as well as cyanobacterial lichenized fungi can show considerable photobiont variability and can have more than one photobiont and even combine green alga and cyanobacteria (Fernandez-Mendoza et al. 2011; Henskens et al. 2012; Nelsen and Gargas 2009; Otalora et al. 2010; Ruprecht et al. 2014; Wornik and Grube 2010). Low photobiont specificity and a high ability to accept different photobionts might be a survival strategy and extend the ecological range of lichens (Blaha et al. 2006; Dal Grande et al. 2017; Leavitt et al. 2015; Ruprecht et al. 2012a; Wirtz et al. 2003). Furthermore, photobiont selection appears to be influenced by abiotic factors like climate (Beck et al. 2002; Fernandez-Mendoza et al. 2011; Peksa and Skaloud 2011; Yahr et al. 2006). At the local scale (for instance along elevational gradients), this may translate into habitat-specific photobiont switches (Vargas Castillo and Beck 2012). Above all, temperature has often been identified as a key factor of photobiont selection of lichens in Antarctica (Green 2009; Kappen and Valladares 2007; Ruprecht et al. 2012a). In warmer regions, myco-/photobiont interactions show increased specificity leading to one-to-one interactions in contrast to more generalist interactions in colder environments (Singh et al. 2017). Thus, it appears that symbiotic interactions in lichens can react very sensitively to environmental change although this conclusion is based on a small database, and these responses have been investigated only in a few species (Allen and Lendemer 2016; Colesie et al. 2014b; Sancho et al. 2017). In general, there is agreement that climatic changes will influence the diversity, abundance and growth of lichens (Sancho et al. 2017) and that lichens therefore represent excellent bioindicators for processes associated with global warming (Alatalo et al. 2015; Allen and Lendemer 2016; Bassler et al. 2016; Sancho et al. 2019).

Over the last decades studies on elevational gradients have re-emerged because the species composition changes remarkably with elevation suggesting a species-specific adaptation to different environmental conditions (Grytnes et al. 2006). They provide steep ecological transitions (e.g. in temperature, humidity and UV radiation) over short distances (Keller et al. 2013; Körner 2007) and several studies suggest that the structure and diversity of communities, the abundance and distribution of species and ecosystem properties and processes can change along elevational gradients (Bassler et al. 2016; Dal Grande et al. 2017; Grytnes et al. 2006; Junker and Larue-Kontic 2018; Körner 2003; Wolf 1993). For lichens, elevational gradients are reported to show large changes in species composition (Dal Grande et al. 2017; Leavitt et al. 2015), habitat-specific photobiont switches (Vargas Castillo and Beck 2012), and/or microclimatic partitioning of ecologically differentiated fungal and algal gene pools (Nadyeina et al. 2014).

This study focuses on saxicolous crustose lichens in continental Antarctica which are associated with green micro alga as photobionts. In general, these lichens are slow growing and restricted to microhabitats on rock surfaces (Hertel 1998), but nevertheless, due to their poikilohydric nature, they are well adapted to habitats with high insolation and with rapid fluctuations in temperature and water availability (Green et al. 2002; Lange 1997; Lange 2000; Schroeter et al. 2011). Most of the lichens analyzed here belong to the ‘lecideoid’ lichen group (Hertel 1984) and these species are assigned to the generic name *Lecidea sensu* Zahlbruckner (1925) but they do not necessarily belong to the genus in its strict sense. Due to their inconspicous growth form, distinguishable only by a few small morphological traits such as spore size and ascus-type, the identification of these lichens is difficult even under best growing conditions (Ruprecht et al. 2019). Extreme climate conditions in cold deserts result in reduced development of the thallus such as chasmolithic growth, a lack of ascomata or sparsely developed ascospores, all features that hamper identification and even the detection of the specimens in the vast landscape (Hertel 2009; Ruprecht et al. 2010). Nevertheless, these pioneers on rocks and pebbles (Hertel 1984; Hertel 1987) belong to one of the most abundant species groups in continental Antarctica (Hertel 2007; Ruprecht et al. 2012b; Ruprecht et al. 2010) and are, therefore, an excellent study system to investiagte changes in symbiotic associations along gradients. The present study covers lecideoid lichen species of the genera *Carbonea* Hertel*, Lecanora* Ach., *Lecidella* Körb., *Rhizoplaca* Zopf and the genus *Lecidea* Ach s.str. (Hertel 1984) plus, in addition, lichen samples of the genera *Austrolecia* Hertel, *Buellia* De Not. and *Huea* C.W. Dodge & G.E. Baker which were included, because they are often have a similar appeareance.

The aim of this study was to analyze the spatial pattern and factors that might affect the distribution of both symbiotic partners within the MDV. We addressed the following objectives: 1) to confirm and extend our knowledge of the abundances of the myco- and photobiont species that have been found by previous, less extensive studies, 2) to investigate the variability of mycobiont/photobiont interactions, in particular, analyze the level of selectivity by using network statistics and 3) to study for mycobiont, photobiont and lichens the relationships with abiotic and climatic factors such as elevation and water status.

## Materials & Methods

### Study area and sampling sites

This study was conducted as part of the New Zealand Terrestrial Antarctic Biocomplexity Survey (nzTABS, http://nztabs.ictar.aq), which was initiated during the International Polar Year 2007–2008, and drew a diverse range of international expertise to profile the biology, geochemistry, geology and climate of the MDV. The study is among the most comprehensive landscape-scale biodiversity surveys undertaken and includes nearly all trophic components found in the MDV ecosystem (Lee et al. 2019). Sampling of soils and biological communities was carried out over two successive Austral summers (2009/10, 2010/11). The geographic area within which lichen samples were collected was the southern part of the MDV (total area: 22700 km^2^, ice-free area: 4500 km^2^; Levy 2013; Fig. 1a, b). The landscape is a mosaic of glacially formed valleys with intervening high ground, ice-covered lakes, ephemeral streams, arid rocky soils, ice-cemented soils, and surrounding glaciers along the steep scree and boulder slopes (Fig. 1b-d; Doran et al. 2002; Stichbury et al. 2011; Yung et al. 2014). There are four main valleys (Miers Valley, Garwood Valley, Hidden Valley and Marshall Valley) and some other extensive ice-free areas (Shangri-La). The topography ranges from sea level to more than 2000 m a.s.l. with granite being the dominant rock type on the ridges and hills, whilst the valley floors are covered with glacial drift. The valleys have the typical glaciated form with a U-cross-section with steep sides, often with scree slopes, which reach up to around 600 m in height. To the west the valleys are separated from the polar plateau by the Royal Society Range that has peaks over 4000 m a.s.l. in height. To the east the valleys open out onto the Ross Ice Shelf, which represents a climatically maritime influenced location within the MDV, despite the absence of ice-free sea at any time of the year (Yung et al. 2014).

**Figure 1:**
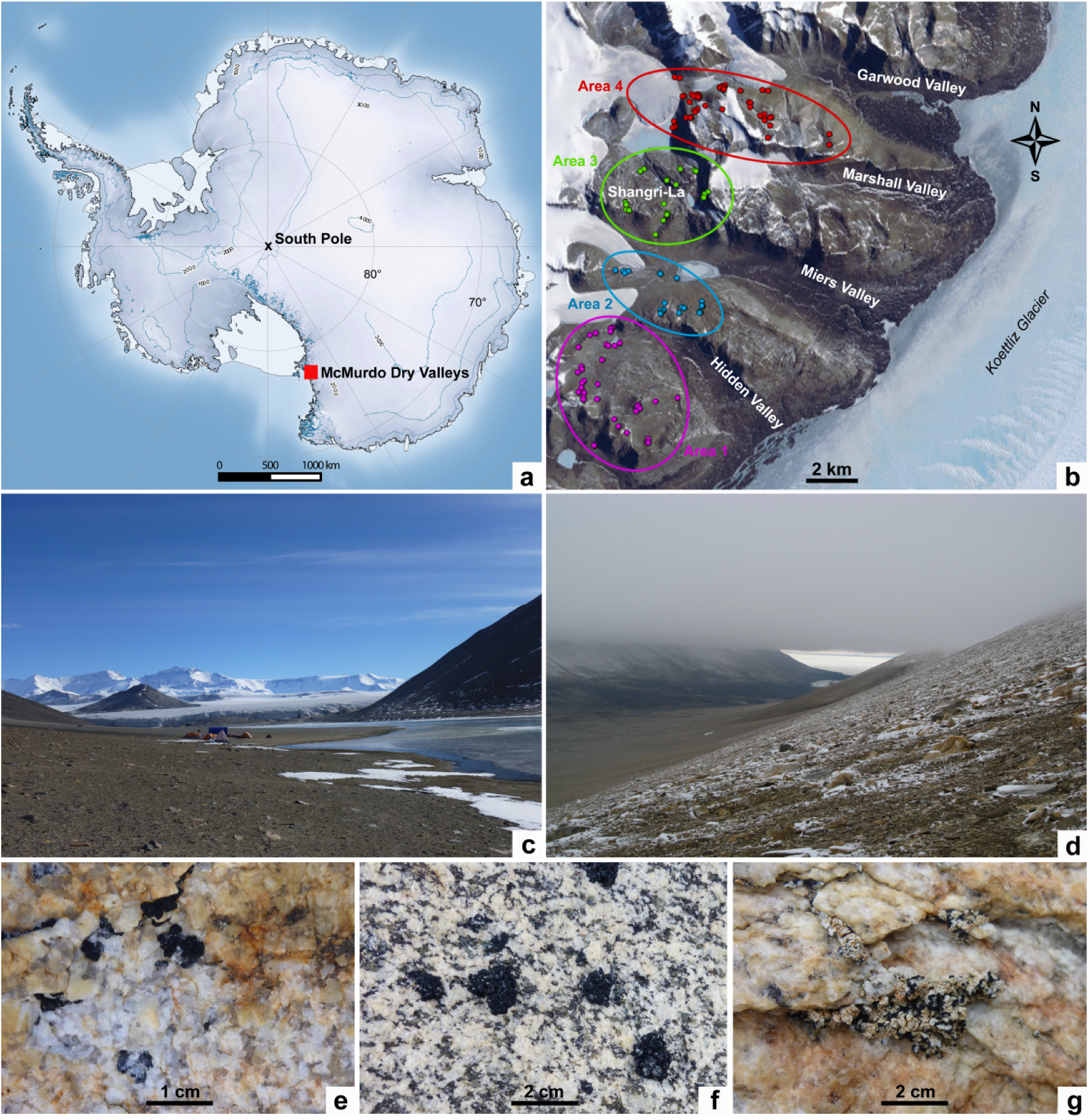
(a) Antarctic continent; investigated area marked with red rectangle (Natural Earth, Qgis), (b) MDV sampling sites defined in four areas (http://www.gpsvisualizer.com), (c) Garwood Valley and field camp, (d) Garwood Valley with incoming cloud bank, (e) *Lecidea cancriformis*, (f) *Rhizoplaca macleanii*, (g) *Austrolecia* sp. 1.

### Climate of the MDV

The climate of the MDV is, for several reasons, classified as that of a polar desert. First, the mountains at the west are sufficiently high to block seaward flowing ice from the East Antarctic ice sheet from reaching the Ross Sea. In addition, the Transantarctic Mountains provide a precipitation shadow, causing an extremely low humidity and lack of snow or ice cover in the MDV (Monaghan et al. 2005). Annual precipitation is < 50 mm a^−1^ water equivalent, with precipitation decreasing away from the coast (Fountain et al. 2010). The major source of liquid water is the seasonal melting of perennial snowbanks and glaciers (Head and Marchant 2014; Stichbury et al. 2011) but, in most cases, this water is not available for lichens that inhabit rock surfaces above the surrounding ground level. MDV climate is best known from the northern valleys, particularly Taylor Valley, because of the McMurdo Long Term Ecological Research programme (McMurdo LTER) that has been active since 1993 (http://www.mcmlter.org). The valley floors of the MDV show mean annual temperatures that range from −30 °C to −15 °C and typically have fewer than 50 days/a where average temperatures exceed 0 °C (Colesie et al. 2014b; Doran et al. 2002; Ochyra et al. 2008). There is agreement that the air temperature lapse rate is close to 1 °C decline per 100 m elevation rise, as well as an increase with distance from the coast to the inland of 0.09 °C per 1 km (McKay 2015). The aspect of the valley slopes has an important impact and north facing slopes are warmer and dryer, south facing slopes are cooler and wetter (Yung et al. 2014). The wind regime is strongly topographically channeled and directed mainly up-or down-valley. During summer, easterly valley winds dominate, due to differential surface heating between the low albedo valley floors and the high albedo ice to the east (Mckendry and Lewthwaite 1990). In winter, wind direction is typically more variable. Cold air pools associated with light winds and very low minimum temperatures (−50 °C) often occupy topographic low points of the valleys during winter (Doran et al. 2002).

Almost all climate information comes from studies on the valley floors. There are, however, conditions that tend to produce a major difference in water regime between valley floors and intervening mountain ranges. First, there is a tendency at higher elevations for greater snowfall and higher humidity, as shown by the presence of clouds at higher elevations (Fig. 1d). Second, there is the regular occurrence within the valleys of what have traditionally been regarded as katabatic winds (Ayling and McGowan 2006; Mckendry and Lewthwaite 1990) but which are now suggested to be foehn winds albeit generated in a slightly different manner to the classic northern hemisphere foehns (Speirs et al. 2010). In the Taylor Valley, for example, these winds are easily recognizable by their sudden arrival, high intensity (around 15 m s^−1^), rapidly rising temperature (by around 25 °C to reach about 0 °C), and rapidly falling relative air humidity to around 20 % (Speirs et al. 2010). These foehn winds also occur in the southern valleys with an example from Miers Valley (Online Resource 2a) showing almost identical characteristics to those in the Taylor Valley. Foehn winds are extremely drying with air vapor pressure deficit rising about 50 times from 0.01 kPa (−30 °C, 80 % RH) to 0.49 kPa (0 °C, 20 % RH). They are also topographically constrained within the valleys and can apparently reach altitudes up to almost 500 m (Speirs et al. 2010). The net result of the higher elevation cold, moister air, and the extremely drying foehn winds within the valleys is that the wetness availability gradient is strongly non-linear and, for the purposes of our analyses, we defined an elevational threshold of about 600 m a.s.l. which marks the upper limit of the steeper valley sides.

### Sample sources

The present study includes 232 lichen samples (lecideoid lichen species of the genera *Carbonea, Lecanora*, *Lecidella*, *Rhizoplaca, Lecidea* and additionally *Austrolecia*, *Buellia* and *Huea*, which have a similar appearence under the extreme climate conditions) collected in four different areas (Hidden Valley, Miers Valley, Shangri La and Garwood Valley) at 153 different localities with a range of aspects (N-facing slopes, flat areas and plateaus) and elevations (171-959 m a.s.l.; Fig. 1a - g, Table 1). 154 specimens were collected between the years 2009 to 2011 by Ulrike Ruprecht and Roman Türk and are deposited at the herbarium of the University of Salzburg (SZU, Online Resource 1a). An additional 78 lichen samples from the same area were obtained from the study of Perez-Ortega et al. (2012) excluding specimens of the genera *Acarospora*, *Caloplaca, Polysporina, Sarcogyne* and *Umbilicaria* (see Online Resource 1b).

**Table 1:** Site descriptions and specifics of the four regions defined within the MDV, including the range of the coordinates of the sampling sites and areas, minimum, maximum and mean values of the elevation of the sampling sites, and the number of lichen samples per area.

|  | Sampling area | Range of coordinates of sampling sites | Min., max. & mean of elevation of sampling sites (m a.s.l.) | Number of sampling sites | Number of lichen samples |
| --- | --- | --- | --- | --- | --- |
| **Area 1** | Hidden Valley | S 78.12° - 78.17°<br>E 163.62° - 163.81° | 361, 645, 911 | 39 | 53 |
| **Area 2** | Miers Lake | S 78.10° - 78.11°<br>E 163.69° - 163.86 | 171, 477, 597 | 28 | 42 |
| **Area 3** | Shangri-La | S 78.05° - 78.08°<br>E 163.71° - 163.87° | 411, 656, 890 | 30 | 51 |
| **Area 4** | Garwood Valley | S 78.02° - 78.05°<br>E 163.80° - 164.10° | 319, 593, 959 | 56 | 86 |

Please note that for most of the data evaluations, mycobionts and photobionts were treated separately. In some analyses (noted in text), only mycobiont species with n ≥ 10 (*min10MycoSp*) were used, whilst others included only photobiont haplotypes with n ≥ 10 (*min10PhoHap*).

### DNA-amplification, purification and sequencing

Total DNA was extracted from the thallus and/or apothecia by using the DNeasy Plant Mini Kit (Qiagen) following the manufacturer’s instructions. For all samples, we sequenced and amplified the internal transcribed spacer (ITS) region of the mycobionts’ and photobionts’ nuclear ribosomal DNA (nrITS). We also amplified additional markers: for the mycobionts the mitochondrial small subunit (mtSSU) and the low-copy protein coding marker RPB1 and, for the photobionts, the chloroplast-encoded intergenic spacer (psbJ-L) and part of the cytochrome oxidase subunit 2 gene (COX2). This was done using newly developed specific primers and PCR-protocols in our project-framework (Ruprecht et al. 2019).

For amplifying nrITS of the mycobiont we used the primers ITS1 (White et al. 1990), ITS1F (Gardes and Bruns 1993), ITS1L (Ruprecht et al. 2019), ITS4 (White et al. 1990), ITS4L (Ruprecht et al. 2019) and for the photobiont 18S-ITS uni-for (Ruprecht et al. 2012a), ITS1T (Kroken and Taylor 2000), ITS1aT (Ruprecht et al. 2014), ITS4T (Kroken and Taylor 2000) and ITS4bT (Ruprecht et al. 2014). For the marker mtSSU the primers CU6 (https://nature.berkeley.edu/brunslab/tour/primers.html), mrSSU1 (Zoller et al. 1999), mtSSU for2 (Ruprecht et al. 2010) and mtSSU rev2 (Ruprecht et al. 2010) and for RPB1, fRPB1-C rev (Matheny et al. 2002), gRPB1-A for (Matheny et al. 2002) and RPB1_for_Lec (Ruprecht et al. 2019) were chosen. For the marker COX2, COXIIf2 and COXIIr (Lindgren et al. 2014) and COX sense (Ruprecht et al. 2019) and for psbJ-L, psbF (Werth and Sork 2010), psbL-sense and psbJ-antisense (Ruprecht et al. 2014) were used. All reactions were performed as described in (Ruprecht et al. 2019). Unpurified PCR-products were sent to Eurofins Genomics/Germany for sequencing.

### Phylogenetic analysis

The sequences of the different marker regions listed above were assembled and edited using Geneious version 6.1.8 (Kearse et al. 2012) and aligned with MAFFTv7.017 (Katoh et al. 2002) for both symbionts. For the photobiont, the classification and labeling of the different operational taxonomical units (OTUs) followed the concept of Leavitt et al. (2015), reevaluated by Ruprecht et al. (2019).

Phylogenetic relationships of the samples of the present study were calculated from the sequences of the marker nrITS. The other makers could not provide further intraspecific variation and were not available for every specimen; therefore they were excluded in all following analyses using sequence data.

A maximum likelihood analysis was calculated with the IQ-TREE web server (Trifinopoulos et al. 2016), using the model selection algorithm ModelFinder (Kalyaanamoorthy et al. 2017). The BIC (Bayesian information criterion) selected for the best-fit model for the mycobiont alignment TN+I+G4 and for the photobiont K2P+I. Branch supports were obtained with the implemented ultrafast bootstrap (UFBoot) (Minh et al. 2013) (number of bootstrap alignments: 1000, maximum iteration: 1000, minimum correlation coefficient: 0.99). Additionally, a SH-aLRT branch test (Guindon et al. 2010) was performed. Each branch of the resulting tree was assigned with SH-aLRT as well as UFBoot supports; the branches with SH-aLRT < 80 % and/ or UFboot < 95 % were collapsed by adding the command *-minsupnew 80/95* to the script.

### Haplotype analysis

We determined the haplotypes (h) of the different mycobiont species and photobiont OTUs by using the function haplotype() of the R package pegas (Paradis 2010) (note: the function only takes into account transversions and transitions but ignores insertions and deletions). For *min10MycoSp* species and photobiont OTUs with h ≥ 2 and at least two haplotypes with n ≥ 3 (*Lecidea cancriformis, Lecidella greenii, Rhizoplaca macleanii* and photobiont OTU *Tr_*A02), haplotype networks were computed, using the function haploNet() of the R package pegas (Paradis 2010). The frequencies were clustered in 10% ranges, for example the circles of all haplotypes making up between 20-30 % have the same size.

### Analysis of spatial distribution

To analyze how the distribution of the lichen specimens correlated with abiotic factors, the sampling sites of the different lichen species or haplotypes in the investigated areas were compared with respect to their environmental specifics. For this we tested the only relevant variable which was elevation. All other variables such as latitude, longitude, and the BIOCLIM variables generated by Wagner et al. (2017) providing a spatial resolution of 1 km, were not suitable for the relatively small area (data not shown). To assure a minimum group size of 10 sample points, the tests only included the *min10MycoSp* species and *min10PhoHap* haplotypes.

In addition, the elevation of the sample sites of the two most dominant photobiont OTUs (*Tr_*A02 and *Tr_*S15) were compared by conducting a nonparametric t-test, using the R function npar.t.test() of the package nparcomp (Konietschke et al. 2015). We used nonparametric multiple comparisons for relative effects (mctp-test; function mctp() of the R package nparcomp (Konietschke et al. 2015), which conducts pairwise comparisons of all possible combinations.

### Analysis of mycobiont – photobiont associations

The associations between mycobiont and photobiont haplotypes were analyzed by computing bipartite networks, using the R function plotweb() of the package bipartite (Dormann et al. 2008). For the bipartite network including mycobiont species and photobiont haplotypes the indices *H*_*2*_’ and *d’* (Blüthgen et al. 2006) were calculated. Both indices are derived from Shannon entropy. *H_2_’* characterizes the degree of complementary specialization or partitioning among the two parties of the entire bipartite network, while *d’* describes the degree of complementary specialization at species or haplotype level. They both range from 0 for the most generalized to 1 for the most specialized case and were computed using the R functions H2fun() and dfun() of the package bipartite (Dormann et al. 2008).

Phylogenetic species diversity of the interaction partners was quantified by calculating a number of further metrices listed in Table 2, including the indices NRI (Net relatedness index), PSV (Phylogenetic species variability) and PSR (Phylogenetic species richness).

**Table 2:** Diversity metrics compared in this study, citations and descriptions of each, and the used R functions.

| Metric | Definition | References | Description and interpretation of values | R function, R package |
| --- | --- | --- | --- | --- |
| $d'$ | Specialization index | (Blüthgen et al. 2006) | Degree of interaction specialization at species level (values range from 0 = most generalized case to 1 = most specialized case) | `dfun()`, `bipartite` (Dormann et al. 2008) |
| NRI | Net relatedness index | (Webb 2000; Webb et al. 2002) | Comparison of phylogenetic distances among all members of a community (pos. values = phylogenetic clustering; neg. values = phylogenetic evenness) | `ses.mpd()`, `picante` (Kembel et al. 2010) |
| PSV | Phylogenetic species variability | (Helmus et al. 2007) | Degree to which species in a community are phylogenetically related (values range from 0 = increased relatedness to 1 = decreased relatedness) | `psd()`, `picante` (Kembel et al. 2010) |
| PSR | Phylogenetic species richness | (Helmus et al. 2007) | PSV multiplied by species richness SR (number of species in a sample); SR after discounting species relatedness (values range from 0 = increased relatedness to SR = decreased relatedness) | `psd()`, `picante` (Kembel et al. 2010) |

### Analysis of DNA polymorphism

For each identified mycobiont and photobiont species with more than one sample, we calculated the haplotype as well as the nucleotide diversity using p-distances with DnaSP v5 (Librado and Rozas 2009). Gaps and missing data were excluded. We focused on *h / N* (number of haplotypes, *h*, divided by number of samples, *N*, per species), *Hd* (haplotype diversity, the probability that two randomly chosen haplotypes are different; Nei 1987) and *π* (nucleotide diversity, average number of nucleotide differences per site between two randomly chosen DNA sequences; Nei and Li 1979).

In order to analyze the dependence of haplotype and nucleotide diversity values on elevation, we used the defined threshold of 600 m a.s.l. The *h / N*, *Hd*, *d’*, and *PSV* of those *min10MycoSp* species with mean values above this threshold were grouped together and then compared to those species with mean values below 600 m a.s.l., using the R function nonpartest() of the package npmv (Ellis et al. 2017), which performs nonparametric comparisons of multivariate samples. *(Note:* π, NRI, *and* PSR *were excluded because of high correlations (r ≥ 0.85) with* h / N (π), Hd (π), d’ (π) *and* PSV (NRI *and* PSR).

## Results

### Phylogenetic analysis

The molecular phylogenies for the mycobiont (Online Resource 2b) and the photobiont (Online Resource 2c) are based on the marker nrITS, because the additional markers (mycobiont: mtSSU, RPB1; photobiont: psbJ-L, COX2) showed little sequence variation in this area. Both analyses include only accessions from the study sites (Online Resource 1a, b) to present the various species- and diversity levels.

#### Mycobiont

The final data matrix for the phylogeny comprised 232 single sequences of the marker nrITS with a length of 610 bp and included sequences of the families *Lecanoraceae, Teloschistaceae, Catillariaceae, Caliciaceae* and *Lecideaceae*. The phylogenetic tree was midpoint rooted and shows a total of 25 strongly supported clades on species level, assigned to eight genera. The backbone is not supported and therefore the topology will not be discussed. The genera *Huea, Austrolecia*, *Buellia* and *Lecidea* are clearly assigned to their family level and are strongly supported. The genera *Carbonea, Lecidella* and *Rhizoplaca* assigned to the family *Lecanoraceae* each form highly supported clades, but do not form an independent clade. The clade of the genus *Lecanora* is divided in five species (*L.* cf*. mons-nivis, L. fuscobrunnea, L.* sp. 1*, L.* sp. 2 *and L.* sp. 3), *Carbonea* in three species (*C. vorticosa*, *C.* sp. URm1 and *C.* sp. 2), *Austrolecia* in three species (*A.* sp. 1, *A.* sp. 2 and *A.* sp. 4), *Buellia* in four species (*B. frigida*, *B.* sp. 1, *B.* sp. 2 and *B.* sp. 3) and *Lecidea* in seven species (*L. andersonii, L. cancriformis, L. lapicida, L. polypycnidophora*, *L.* sp. 5, *L.* sp. 6 and *L.* UCR1). The samples allocated to the genera *Lecidella*, *Rhizoplaca* and *Huea* were, in each genus, monospecific (*Lecidella greenii*, *Rhizoplaca macleanii* and *Huea sp*. 1). The taxonomical assignment of the obtained sequences was made using the following literature: Perez-Ortega et al. (2012); Ruprecht et al. (2012b); Ruprecht et al. (2010).

#### Photobiont

The final data matrix for the phylogeny comprised 222 single sequences of the marker nrITS with a length of 578 bp. The phylogenetic tree was midpoint rooted and shows four strongly-supported clades, all of them belonging to the genus *Trebouxia*. They were assigned to OTU level (Puillandre et al. 2012) using the system of Leavitt et al. (2015), reevaluated by (Ruprecht et al. 2019) and assigned to *Tr_*A02, *Tr_*S02, *Tr_*S15, *Tr_*S18. The OTU *Tr_*A02 was by far the most common, being present in 202 (91 %) of our 222 samples. Photobiont sequences taken from Perez-Ortega et al. (2012) which were labelled only with numbers were included in our system of assigning the haplotypes to the appropriate OTUs (Ruprecht et al. 2019) and therefore renamed.

### Haplotype analysis

The following analyses were based on the nrITS sequences of myco- and photobionts. The number of haplotypes differed significantly between myco- and photobionts. We identified 48 different mycobiont but only 17 different photobiont haplotypes. The most frequent mycobiont haplotype was *Lecidella greenii*_h01 with 28 samples, the most frequent photobiont haplotype was *Tr_*A02_h01 with 87 samples. Additionally, some of the mycobiont species appear to be more diverse (number of different haplotypes) than others. **Fehler! Verweisquelle konnte nicht gefunden werden.** & 3 show bar charts that give the number of samples per mycobiont species/photobiont OTU per area and per haplotype.

Three different mycobiont species (*Lecidea cancriformis*, *Lecidella greenii* and *Rhizoplaca macleanii*) and the most common photobiont OTU (*Tr_*A02) met the required criteria defined above for the construction of haplotype networks (h ≥ 2 and at least two haplotypes with n ≥ 3). In Fig. 4 the respective haplotype networks show the spatial location within the four areas. As shown in Fig. 2 for mycobiont species/photobiont OTU, the distribution again turned out to be rather uniform, with most of the haplotypes found in all of the four areas.

**Figure 2:**
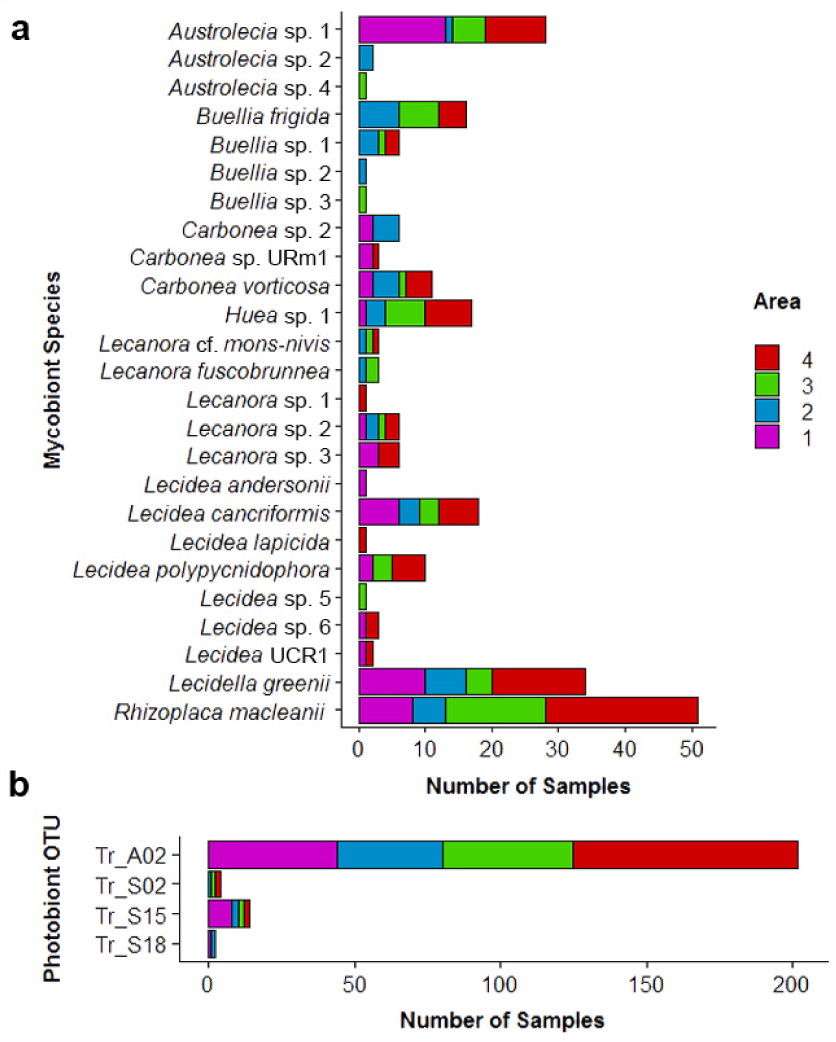
Number of samples per species/OTU and area (cf. Fig. 1b). (a) mycobiont species (total sample size: n = 232), (b) photobiont OTUs (total sample size: n = 222).

**Figure 3:**
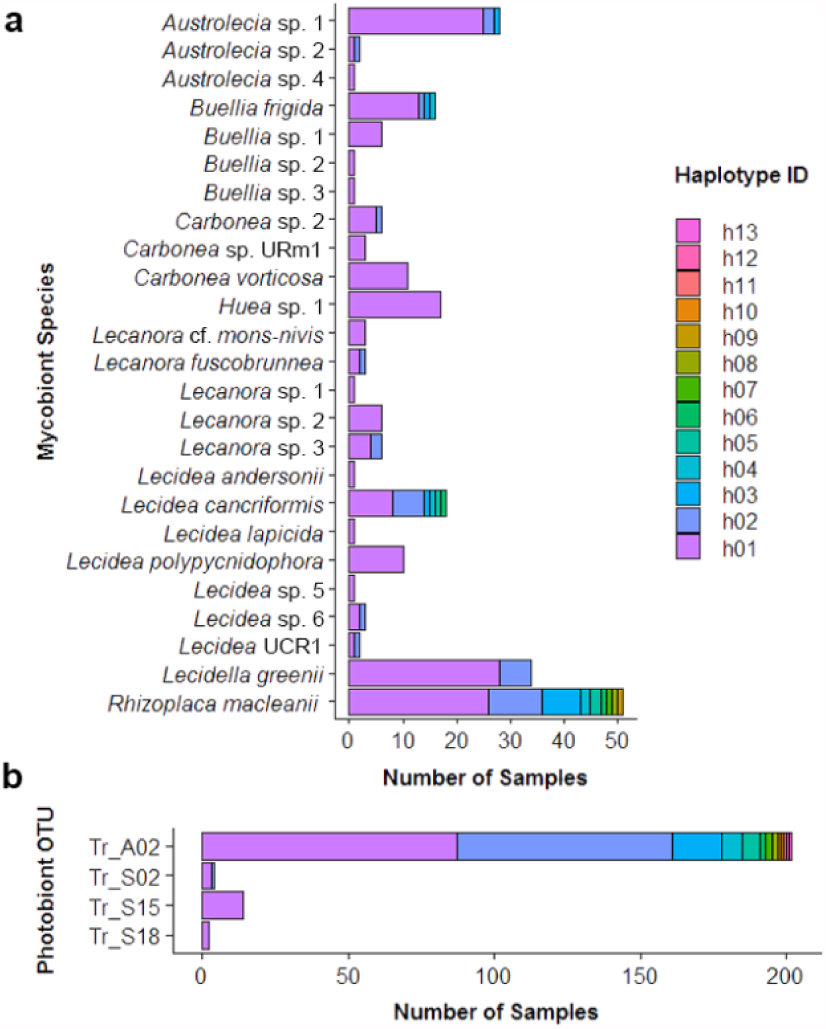
Number of samples per species/OTU and haplotype. (a) mycobiont species (total sample size: n = 232), (b) photobiont OTUs (total sample size: n = 222).

**Figure 4:**
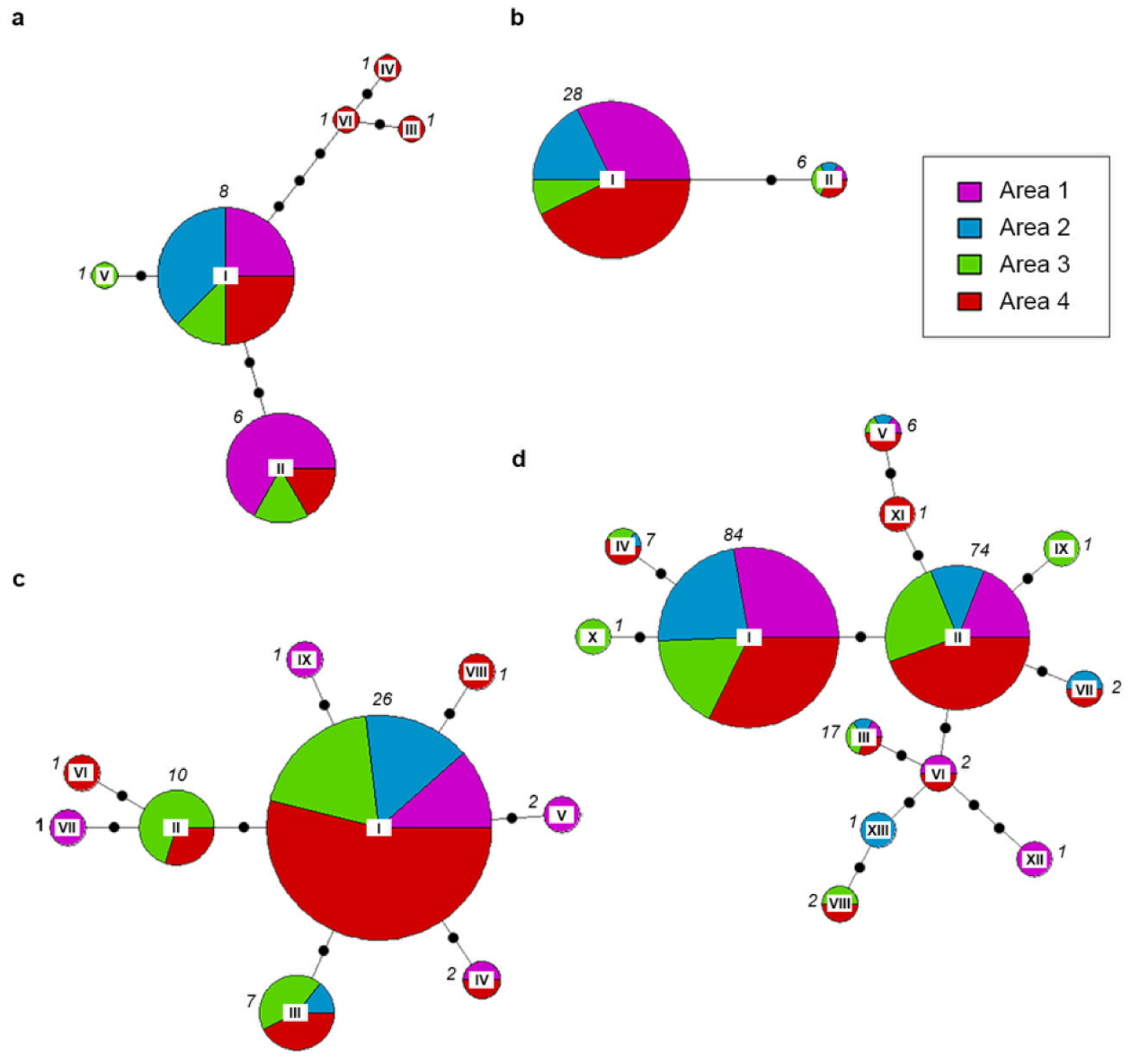
Haplotype networks of those mycobiont species/ photobiont OTUs with h ≥ 2 and at least two haplotypes with n ≥ 3: (a) *Lecidea cancriformis*, (b) *Lecidella greenii*, (c) *Rhizoplaca macleanii* and the photobiont OTU *Tr_*A02, (d) showing the spatial distribution within Area 1 to Area 4 (cf. Fig. 1b & 2). Roman numerals at the center of the pie charts refer to the haplotype IDs, italic numbers next to the pie charts to the total number of samples per haplotype. The circle sizes reflect relative frequency within the species/OTU; in doing so, frequencies were clustered in ten, so that for example the circles of all haplotypes making up between 20-30 % have the same size.

### Analysis of spatial distribution

For 12 of the 28 pairwise comparisons for the mycobionts species (*min10MycoSp*) and photobiont haplotypes (*min10PhoHap*) the mctp-tests (pairwise comparisons of all possible combinations) for elevation showed significant differences, which are also visually recognizable when comparing the maps of the sample locations where the sample locations for each species are shown separately with varying colors indicating their elevations (*min10MycoSp* and *min10PhoHap* summarized in the respective OTUs; Online Resource 2d). The pairs with significant differences as well as the associated p-values are given in Online Resource 1c. For the photobionts the mctp-test showed significant differences for only for two of the six pairwise comparisons (*Tr_*A02_h01 and *Tr_*A02_h03, *Tr_*A02_h01 and *Tr_*S15_h01 and, in each case, *Tr_*A02_h01 has lower elevation values; Online Resource 1c). The nonparametric t-test for comparing the elevation of the sample locations on OTU level resulted in a significant difference between the two groups for *Tr_*A02 and *Tr_*S15 (p = 0.005) with sampling sites of the OTU *Tr_*S15 being higher.

Fig. 5 shows the elevational distribution of the different mycobiont species and photobiont OTUs at their sample sites for *min10MycoSp* species and *min10PhoHap* haplotypes. The mycobiont species *Lecidea cancriformis* and *Rhizoplaca macleanii* had a significant tendency to higher elevations, whilst *Carbonea vorticosa*, *Lecidea polypycnidophora* and *Lecidella greenii* had a significant tendency to lower elevations. One haplotype of the photobiont, OTU *Tr_*S15, was restricted to higher elevations. The remaining mycobiont species and photobiont haplotypes have an intermediate distribution.

**Figure 5:**
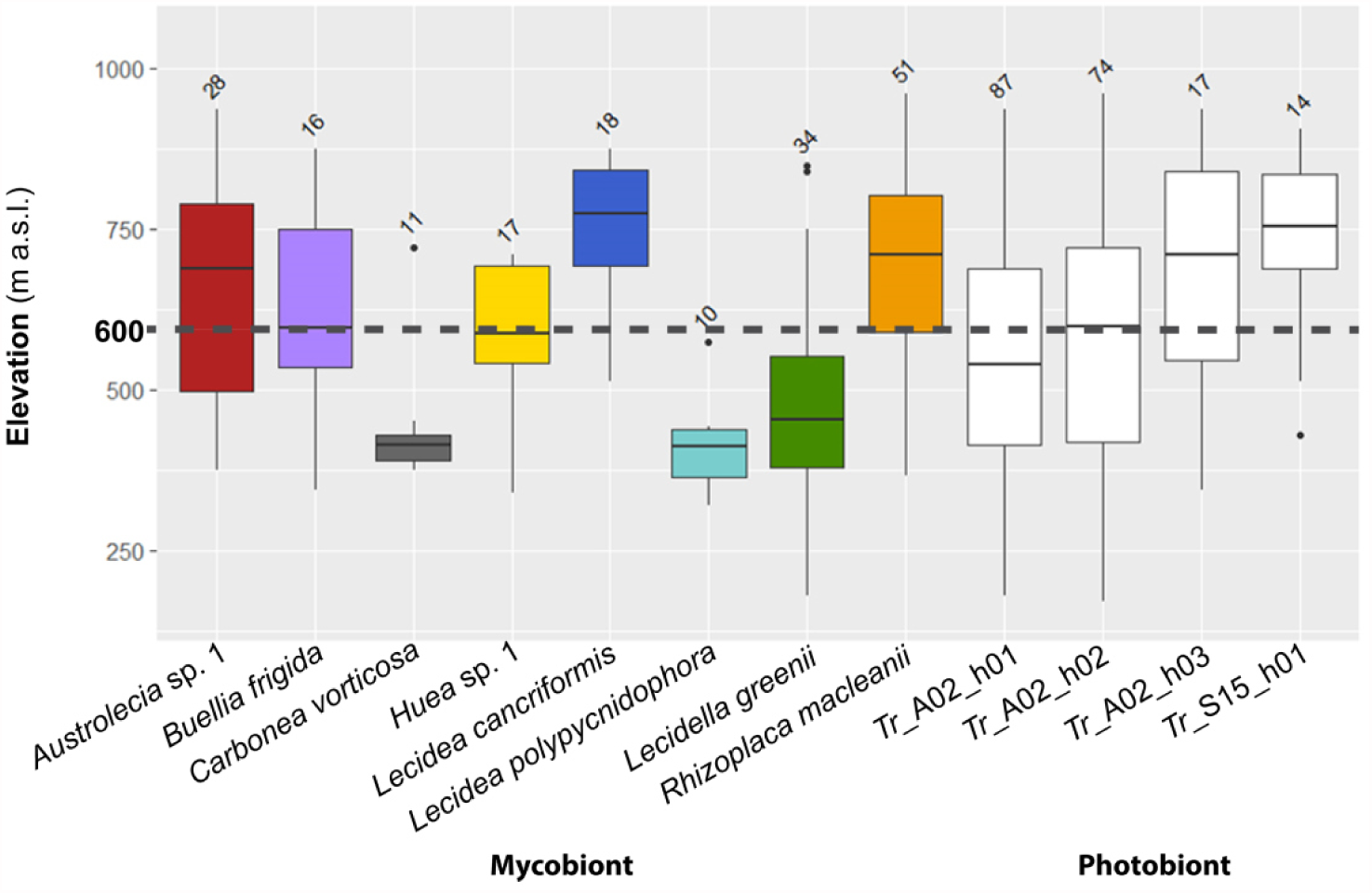
Boxplots showing the elevation of the sample sites of the *min10MycoSp* species and *min10PhoHap* haplotypes. Numbers in italics refer to sample sizes. The elevational threshold of 600 m a.s.l. is highlighted with a dashed line.

### Analysis of mycobiont-photobiont associations

The bipartite network was calculated for all associations between the mycobiont species (*min10MycoSp;* lower level) and the respective photobiont haplotypes (higher level; Fig. 6). The *H_2_’* (overall level of complementary specialization of all interacting species) of this network shows a low value of 0.226 which indicates a network with mostly generalized interactions (as opposed to specialized). This was mainly caused by the dominant occurrence of the three most abundant haplotypes (h01-h03) of the most common photobiont OTU *Tr_*A02 (91 % of 222 accessions). All mycobiont species were additionally associated to a variety of other and less abundant haplotypes (h04-h13) of this OTU *Tr_*A02. Furthermore, some of the mycobionts were associated with accessions of the distantly related OTUs *Tr_*S02, *Tr_*S15 and *Tr_*S18. The network matrix shows the number of associations between the mycobiont species and photobiont haplotypes (Online Resource 1d). The individual *d’* values (complementary specialization at species or haplotype level) are ranging from 0 to 1 and are presented in Table 3.

**Figure 6:**
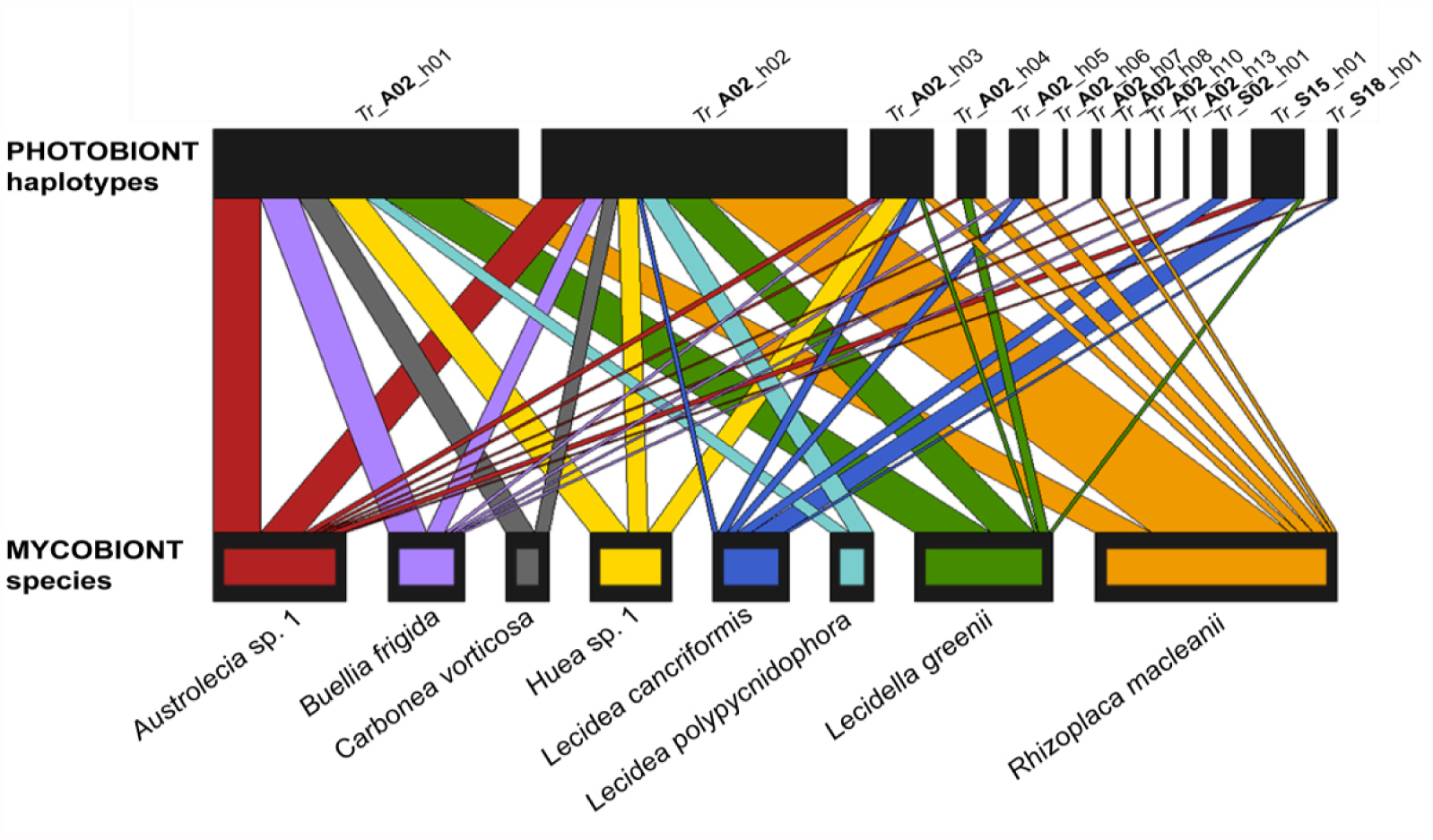
Bipartite network including the *min10MycoSp* species as well as the haplotypes of their associated photobionts with an *H_2_’* = 0.226. Rectangles represent species, and the width is proportional to the number of samples. Associated species are linked by lines, whose width is proportional to the number of associations.

**Table 3:** Diversity (left) and specificity indices (right) for the different mycobiont species and photobiont OTUs: N, number of sequences; h, number of haplotypes; h / N, quotient of *h* and *N;* Hd, haplotype diversity; π, nucleotide diversity; d’, specialization index; NRI, net relatedness index; PSV, phylogenetic species variability; SR, species richness; PSR, phylogenetic species richness.

| Mycobiont Species | N | h | h / N | Hd | $\pi$ | $d'$ | NRI | PSV | SR | PSR |
| --- | --- | --- | --- | --- | --- | --- | --- | --- | --- | --- |
| *Austrolecia* sp. 1 | 28 | 3 | 0.107 | 0.204 | 0.00069 | 0.053 | -0.127 | 0.454 | 8 | 3.636 |
| *Austrolecia* sp. 2 | 2 | 2 | 1.000 | 1.000 | 0.00791 | 0.140 | - | - | 1 | - |
| *Austrolecia* sp. 4 | 1 | 1 | 1.000 | - | - | 0.000 | - | - | 1 | - |
| *Buellia frigida* | 16 | 4 | 0.250 | 0.350 | 0.00098 | 0.105 | 2.210 | 0.029 | 6 | 0.173 |
| *Buellia* sp. 1 | 6 | 1 | 0.167 | 0.000 | 0.00000 | 0.014 | -0.784 | 0.670 | 3 | 2.011 |
| *Buellia* sp. 2 | 1 | 1 | 1.000 | - | - | 0.000 | - | - | 1 | - |
| *Buellia* sp. 3 | 1 | 1 | 1.000 | - | - | 0.036 | - | - | 1 | - |
| *Carbonea* sp. 2 | 6 | 2 | 0.333 | 0.333 | 0.00073 | 0.163 | -0.354 | 0.511 | 4 | 2.046 |
| *Carbonea* URm1 | 3 | 1 | 0.333 | 0.000 | 0.00000 | 0.000 | 0.895 | 0.011 | 2 | 0.023 |
| *Carbonea vorticosa* | 11 | 1 | 0.091 | 0.000 | 0.00000 | 0.064 | 0.895 | 0.011 | 2 | 0.023 |
| *Huea* sp. 1 | 17 | 1 | 0.059 | 0.000 | 0.00000 | 0.120 | 1.266 | 0.023 | 3 | 0.069 |
| *Lecanora* cf. *mons-nivis* | 3 | 1 | 0.333 | 0.000 | 0.00000 | 0.305 | 0.819 | 0.046 | 2 | 0.092 |
| *Lecanora fuscobrunnea* | 3 | 2 | 0.667 | 0.667 | 0.00129 | 0.391 | -1.239 | 1.000 | 2 | 2.000 |
| *Lecanora* sp. 1 | 1 | 1 | 1.000 | - | - | 0.366 | - | - | 1 | - |
| *Lecanora* sp. 2 | 6 | 1 | 0.167 | 0.000 | 0.00000 | 0.048 | 1.285 | 0.015 | 3 | 0.045 |
| *Lecanora* sp. 3 | 6 | 2 | 0.333 | 0.533 | 0.00104 | 0.254 | 1.590 | 0.025 | 4 | 0.099 |
| *Lecidea andersonii* | 1 | 1 | 1.000 | - | - | 0.366 | - | - | 1 | - |
| *Lecidea cancriformis* | 18 | 6 | 0.333 | 0.719 | 0.00388 | 0.595 | -1.273 | 0.665 | 6 | 3.990 |
| *Lecidea lapicida* | 1 | 1 | 1.000 | - | - | 0.000 | - | - | 1 | - |
| *Lecidea polypycnidophora* | 10 | 1 | 0.100 | 0.000 | 0.00000 | 0.059 | 0.895 | 0.011 | 2 | 0.023 |
| *Lecidea* sp. 5 | 1 | 1 | 1.000 | - | - | 1.000 | - | - | 1 | - |
| *Lecidea* sp. 6 | 3 | 2 | 0.667 | 0.667 | 0.01028 | 0.147 | - | - | 1 | - |
| *Lecidea* UCR1 | 2 | 2 | 1.000 | 1.000 | 0.00193 | 0.491 | 0.866 | 0.023 | 2 | 0.045 |
| *Lecidella greenii* | 34 | 2 | 0.059 | 0.299 | 0.00062 | 0.049 | 0.027 | 0.415 | 5 | 2.074 |
| *Rhizoplaca macleanii* | 51 | 9 | 0.176 | 0.692 | 0.00185 | 0.155 | 2.528 | 0.033 | 7 | 0.234 |
| Photobiont OTU | N | h | h / N | Hd | $\pi$ | $d'$ | NRI | PSV | SR | PSR |
| Tr_A02 | 202 | 11 | 0.054 | 0.659 | 0.00174 | 0.367 | 1.235 | 0.485 | 42 | 20.375 |
| Tr_S02 | 4 | 2 | 0.500 | 0.500 | 0.00189 | 0.721 | -0.336 | 0.441 | 2 | 0.882 |
| Tr_S15 | 14 | 1 | 0.071 | 0.000 | 0.00000 | 0.493 | -0.965 | 0.529 | 7 | 3.705 |
| Tr_S18 | 2 | 1 | 0.500 | 0.000 | 0.00000 | 0.344 | -1.188 | 1.000 | 2 | 2.000 |

### Species richness of mycobiont vs photobiont

The number of different symbiotic partners at haplotype level (SR) as a function of the number of mycobiont haplotypes (*h* in Table 3) is illustrated in Fig. 7 for the *min10MycoSp* species. The two variables show a correlation of r = 0.701; thus, highly variable mycobionts tend to be associated with a higher number of photobiont haplotypes.

**Figure 7:**
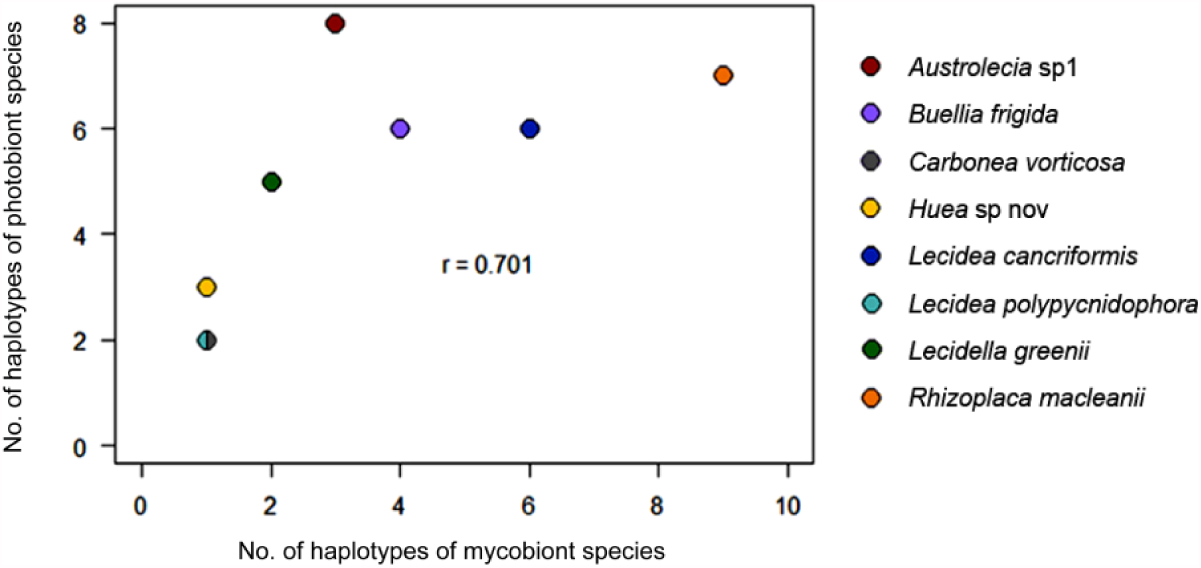
Scatterplot of the species richness (SR) of photobiont haplotypes dependent on the number of haplotypes (*h*) of each *min10MycoSp* species (*Carbonea vorticosa* and *Lecidea polypycnidophora* share the same coordinates) including the Pearson correlation coefficient (*r*).

### Analysis of DNA polymorphism and nonparametric comparisons of multivariate samples

Analyses of DNA polymorphism and nonparametric comparisons of multivariate samples were achieved for *min10MycoSp* and *min10PhoHap* including the parameters *h / N* (number of haplotypes, *h*, divided by number of samples, *N*, per species), *Hd* (haplotype diversity), and *π* (nucleotide diversity), *d’* (specialization index), NRI (net relatedness index), PSV (phylogenetic species variability), SR (species richness) and PSR (phylogenetic species richness; Table 3).

The lowest values of *Hd* and *π* were developed by four mycobiont species of *min10MycoSp* (*C. vorticosa, L. polypycnidophora, L. greenii* and *Huea* sp. 1) which occur at the lowest and intermediate elevations. In contrast, species that were found only at the higher elevations (*R. macleanii, L. cancriformis*) showed the opposite behavior and had the highest values for *Hd* and *π*. These results were supported by nonparametric tests of the different haplotype and nucleotide diversity values which showed a significant difference between *min10MycoSp* samples below the threshold of 600 m a.s.l. (*Carbonea vorticosa*, *Huea* sp. 1, *Lecidea polypycnidophora* and *Lecidella greenii*) and those above (*Austrolecia* sp. 1, *Buellia frigida*, *Lecidea cancriformis* and *Rhizoplaca macleanii*) in terms of *h / N*, *Hd*, *d’* and PSV (ANOVA type test p-value: 0.009). The scatterplots of the indices as a function of elevation as well as the Pearson correlation coefficients are presented in Online Resource 2e.

## Discussion

The McMurdo Dry Valleys at 78° S in Victoria Land, Antarctica are known as the largest continuous ice-free landscape on the continent and are mostly colonized by lithic and soil related microbial communities (Bottos et al. 2014; Colesie et al. 2014a; De Los Rios et al. 2014; Lee et al. 2019; Pérez-Ortega et al. 2012; Yung et al. 2014). The only comprehensive evaluation to date for saxicolous lichens was that of Pérez-Ortega et al. (2012) which showed a high number of mycobiont species (26) and, in contrast, a low number of photobiont species (four). Here we use a new and much larger dataset focused specifically on the lecideoid lichen group that includes some other species with similar morphologies in the same and adjacent areas and the respective data of Perez-Ortega et al. (2012; Fig. 1b). A total of 25 mycobiont species were identified with a composition that was mostly similar to the previous study but including a few additional specimens (*Lecanora fuscobrunnea*, unknown *Lecidea* and *Buellia* species). They were all associated with the same set of common photobionts (four species), dominated (91%) by the more recently reclassified OTU *Trebouxia* A02 (Leavitt et al. 2015), equivalent to the species *T.* sp. URa2 (Ruprecht et al. 2012a). The evaluation of Pérez-Ortega et al. (2012) included not only the same set of lecideoid and morphologically similar lichen genera (*Lecidea; Carbonea, Lecanora, Lecidella, Rhizoplaca; Austrolecia; Buellia*; *Huea*) as investigated in this study, but also five additional lichen genera (*Acarospora; Caloplaca; Polysporina; Sarcogyne; Umbilicaria*).

In our study, we found that the different mycobiont species and photobiont OTUs within the MDV appear to be relatively evenly distributed across all four primary sample sites (Fig. 2), which is in basic agreement to the previous study. However, our results show distinct patterns for distribution, genetic diversity and specificity. These results contrast with Pérez-Ortega et al. (2012) where the distribution of mycobionts and photobionts was independent of elevation and other abiotic factors. A clear trend has now emerged showing that the distribution of species/OTUs is significantly related to elevation, using 600 m a.s.l. as a defined threshold dividing higher and lower sites. The mycobiont species *Carbonea vorticosa*, *Lecidea polypycnidophora* and *Lecidella greenii* were found almost exclusively below and *Lecidea cancriformis* and *Rhizoplaca macleanii* above the threshold (Fig. 5), which was supported by mctp-tests for pairwise comparisons (Online Resource 1c).

In contrast, the dominant photobiont OTU *Tr_*A02 is distributed everywhere whilst the remaining and distantly related OTUs (*Tr_*S02, *Tr_* S15 and *Tr*_ S18) are mostly restricted to the higher elevation habitats (cold and humid; Fig. 5, Online Resource 2c). This result is in agreement with Dal Grande et al. (2018), who reported clear elevational preferences for some *Trebouxia* taxa at the OTU level at a mountain range in central Spain covering an elevational gradient of 1400 m. He suggested that altitude plays a prominent role in shaping the community structure of these algae. The distribution of *Tr*_S15 (equivalent to the species *T.* URa1), only occurring in the cold and humid areas was surprising because the first specimen of this species described was found at the climatically most extreme and dry habitats in the Darwin area 80° S (Ruprecht et al. 2012a). However, lichen photobionts seem to have clear ecological preferences and niche differentiations. This was also shown for various *Asterochloris* lineages (Trebouxiophyceae) which have diverging preferences in terms of rain and sun exposure (Peksa and Skaloud 2011) whilst Nadyeina et al. (2014) reported elevational partitioning in the distribution of different gene pools of the photobiont of the lungwort lichen *Symbiochloris reticulate* (Trebouxiophyceae; Skaloud et al., 2016). In general, several studies suggest a strong genetic association of lichen-associated algae with climatic factors and substrate (e.g. Fernandez-Mendoza et al. 2011; Vargas Castillo and Beck 2012; Yahr et al. 2006), and this has been interpreted as evidence for ecological specialization (Muggia et al. 2014; Ruprecht et al. 2012a).

The mycobiont species *min10MycoSp* not only show clear spatial differentiation with respect to elevation for species and OTUs but also for variables expressing the genetic diversity and specialization towards both symbiotic partners. A higher elevation correlates with a higher number of haplotypes (*Hd*) and an increased nucleotide diversity (*π)* which leads to a greater intraspecific diversity within the mycobionts (Table 3). These differences are also partially reflected by a higher *d’*, PSV, PSR and a low NRI which show a low relatedness to the co-occurring photobiont, associated with the rarely occurring and highly differentiated other OTUs *Tr_*S02, *Tr_*S15 and *Tr_*S18. Consequently, mycobionts with a high genetic diversity have a higher number of interacting partners. These findings are partially supported by the study of Singh et al. (2017), who reported climate as a selective pressure in terms of increased specificity of myco-/photobiont interactions.

Our study has also shown that highly variable mycobionts are associated with a larger number of photobiont haplotypes (Fig. 7). If we focus on the species which are significantly distributed either below or above the threshold of 600 m a.s.l. three main scenarios emerged: (1) mycobionts with low genetic diversity (*Carbonea vorticosa*, *Lecidea polypycnidophora* and *Lecidella greenii*) are associated with one photobiont OTU *Tr*_A02, and were found in only the lower area; (2) a mycobiont with a high genetic diversity (*Rhizoplaca macleanii*) is still solely associated to one photobiont OTU (*Tr*_A02) and is only located at the high elevated areas and (3) the mycobiont with the highest genetic diversity (*Lecidea cancriformis*) is associated with highest number of photobiont OTUs, in the high elevation sites. These findings are in agreement with the known distribution of *L. greenii* and *Tr_*A02 (*Trebouxia* URa2), which, so far, have only been reported for sites in the more northern parts of the Ross Sea region and have never been found at the most extreme southern environments like the Darwin area (Ruprecht et al. 2012a; Ruprecht et al. 2012b). In contrast, *L. cancriformis* is one of the most widespread lichens, being distributed all over Continental Antarctica and is associated with all known photobiont species (Castello 2003; Ruprecht et al. 2012a; Ruprecht et al. 2010).

The above results suggest that the mycobionts are dependent on the availability of climatically adapted photobionts. However, the mycobionts seem to have also their unique climate specific preferences because they do not make use of the whole climate niche of the associated photobionts. These findings are only partially in line with previous studies (e.g. Romeike et al. 2002; Wirtz et al. 2003) that suggest that in extreme environments like the Antarctic continent there might be a selection pressure against photobiont specificity so that more versatile mycobionts are favored. Flexibility concerning the partner choice has been considered as an adaptive strategy to survive harsher environmental conditions (Leavitt et al. 2013; Singh et al. 2017; Werth and Sork 2010).

To explain the myco- and photobiont distribution we need to understand what abiotic factors control terrestrial life in these polar ecosystems. For the MDV Lee et al. (2019) state clearly that one of the most important factors governing the distribution of taxa in the MDV is temperature, which is accepted to be inversely correlated to elevation (McKay 2015). However, much less is known about the conditions for wetness and humidity. The available wetness index for the MDV quantifies the expected wetness of a unit within the watershed by calculating the amount of possible water flowing into that unit from estimated snow fall (Stichbury et al. 2011). For rock associated lichens this is not relevant, because they are not connected to this source of water. They are, therefore, dependent on moisture provided by the very low precipitation, infrequent melting snow (Head and Marchant 2014) and humidity provided by incoming fog and clouds from the sea. Additionally, it is now clear if the occasional foehn wind events cause severe drying within the valleys at altitudes up to 500 m. At higher elevations there is cold and moister air and this establishes a strong moisture availability gradient with elevation (Speirs et al. 2010; Fig. 1d). Our results suggest that our defined elevation threshold of about 600 m a.s.l. is a reasonable level which marks the shift from lower, dryer to higher, more humid conditions. Habitat aspect is also known to be important. Yung et al. (2014) described large differences with respect to just aspect for their chasmoendolithic microbial communities at Miers Valley. Similar results were also reported for the more maritime site, Botany Bay (Seppelt et al. 2010). We did not find any impact of other topographical features such as distance to coast, slope, aspect and substrate. The collection sites were mainly N-facing or on plateaus, our transects were narrow and consistently only five to ten km inland plus the underlying rock in the whole area is granite and the investigated lichens are restricted to siliceous rock (Ruprecht et al. 2012b; Ruprecht et al. 2010). However, our sampling was equally distributed below and above the threshold of 600 m a.s.l., so the differences found for species distribution, genetic diversity and specificity appears to be due to the changing climate conditions, particularly moisture, along the elevational gradient.

## Acknowledgements

We are grateful to T.G.A Green (Waikato, NZ; Madrid, E), M. Affenzeller, A. Blaschka, G. Brunauer, A.M. Zimmermann (Salzburg, A) for valuable advice and R. Türk (Salzburg, A) for collecting one third of the lichen samples in 2010 and providing the picture for Fig. 1d. The lichen collections were supported by grants to SCC from the New Zealand Foundation for Research, Science and Technology (UOWX0710), the New Zealand Ministry of Business, Innovation and Employment (UOWX1401) and the University of Waikato. Antarctica New Zealand is thanked for logistic support. This study and open access was financially supported by the Austrian Science Fund (FWF): P26638_B16, Diversity, ecology and specificity of Antarctic lichens.

## Online Resource 1

**Online Resource 1a**: Samples used in this study, with information on haplotypes, collecting localities and Genbank accession numbers of different markers. To avoid redundant data accumulation, for every haplotype only one reference sequence was uploaded to Genbank.

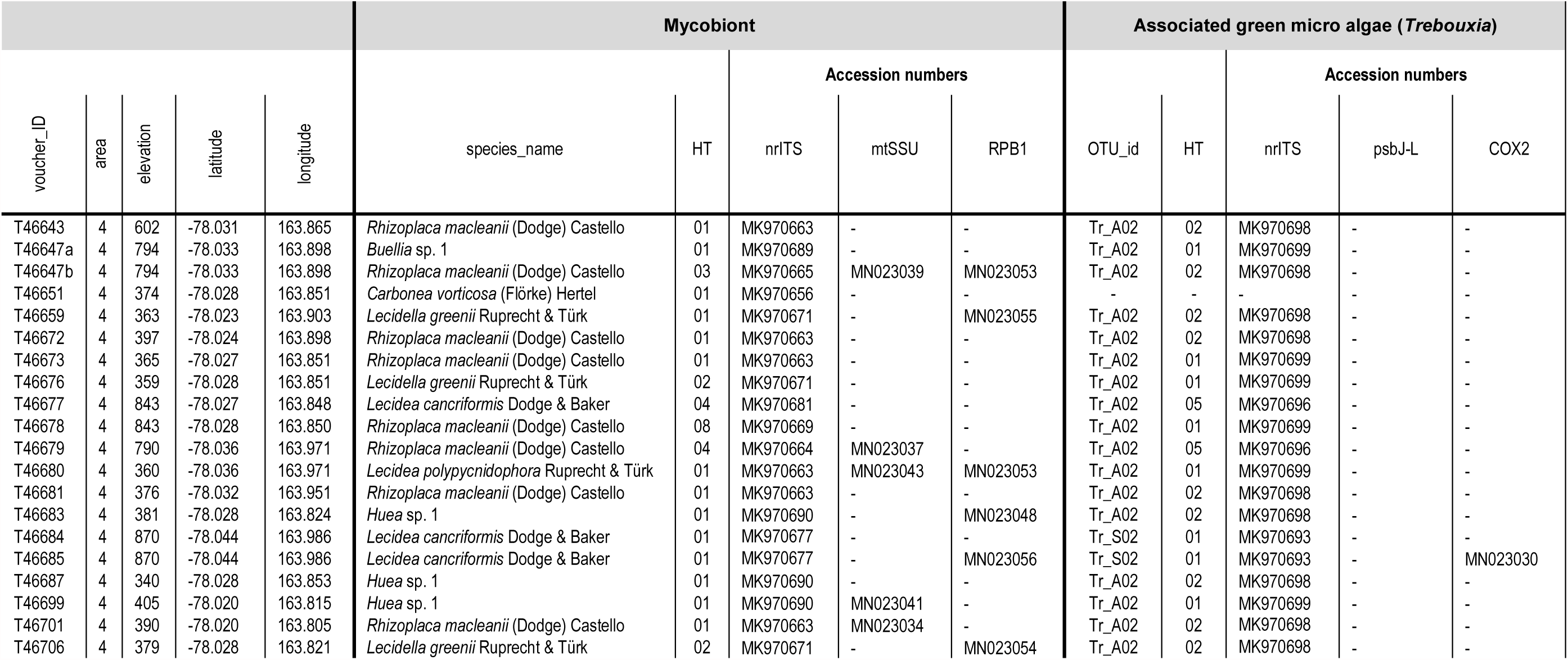
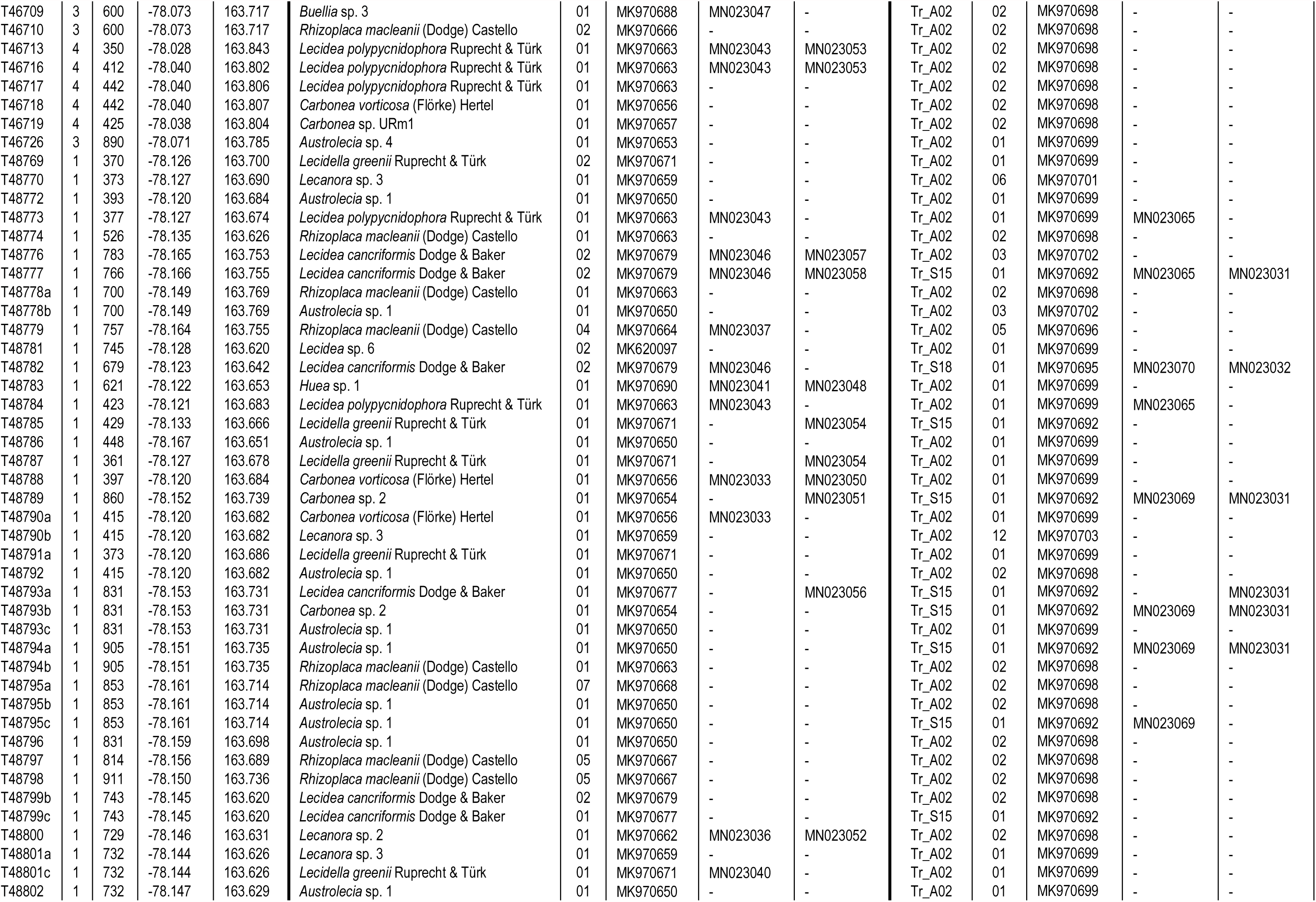
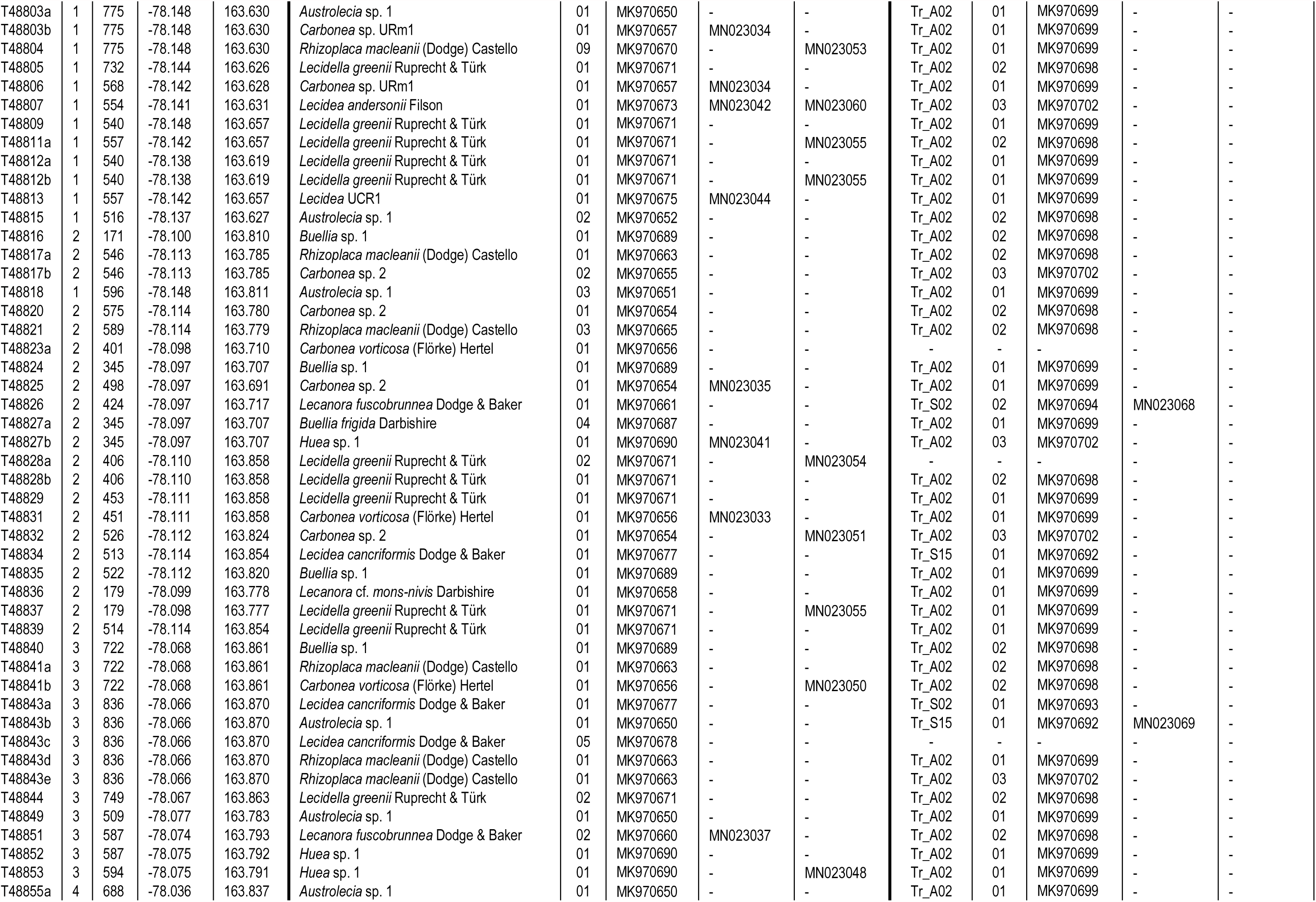
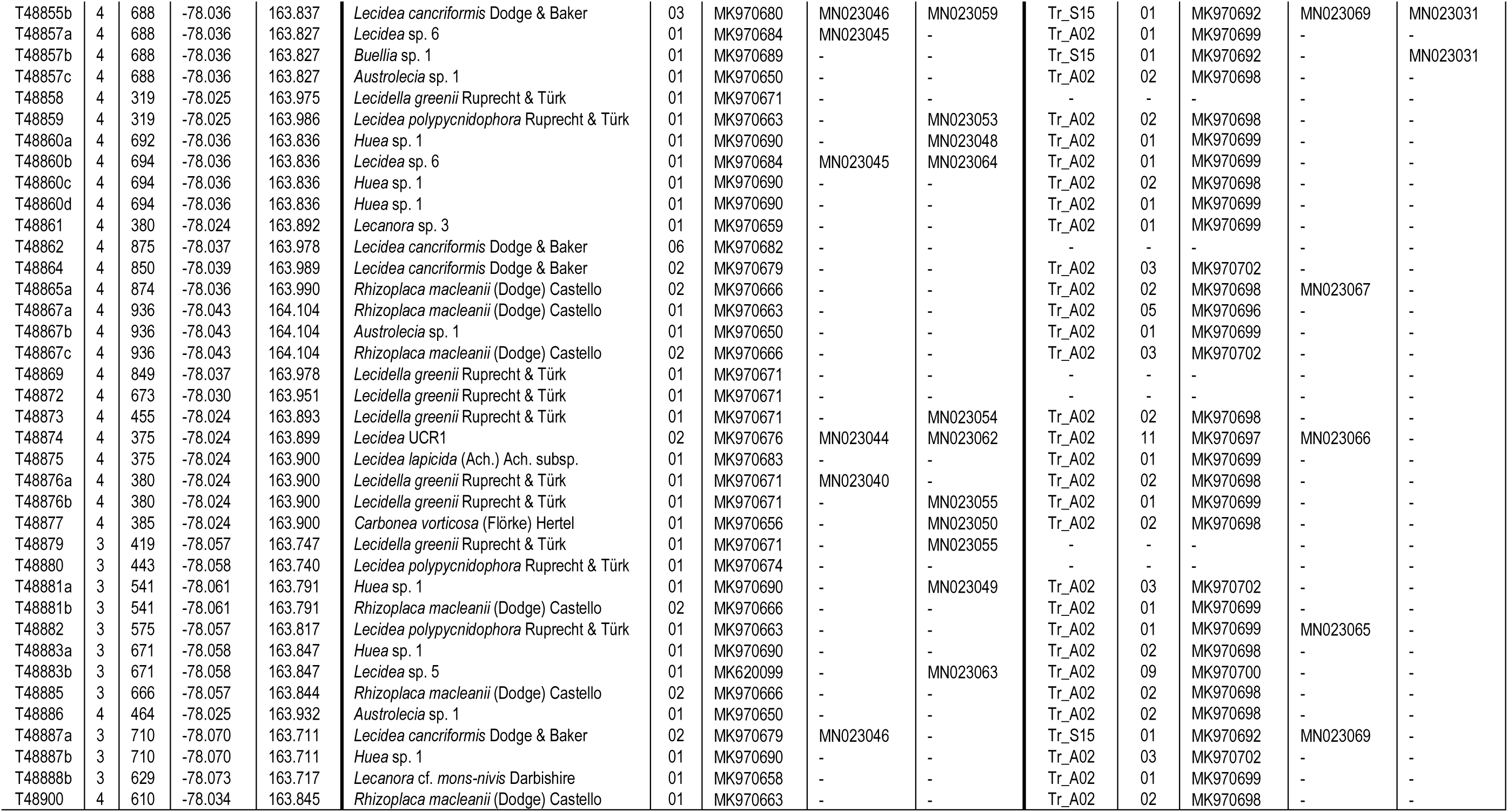

|  |  |  |  |  | Mycobiont |  |  |  |  | Associated green micro algae (*Trebouxia*) |  |  |  |  |
| --- | --- | --- | --- | --- | --- | --- | --- | --- | --- | --- | --- | --- | --- | --- |
| voucher_ID | area | elevation | latitude | longitude | species_name | HT | Accession numbers |  |  | OTU_id | HT | Accession numbers |  |  |
|  |  |  |  |  |  |  | nrITS | mtSSU | RPB1 |  |  | nrITS | psbJ-L | COX2 |
| T46643 | 4 | 602 | -78.031 | 163.865 | *Rhizoplaca macleanii* (Dodge) Castello | 01 | MK970663 | - | - | Tr_A02 | 02 | MK970698 | - | - |
| T46647a | 4 | 794 | -78.033 | 163.898 | *Buellia* sp. 1 | 01 | MK970689 | - | - | Tr_A02 | 01 | MK970699 | - | - |
| T46647b | 4 | 794 | -78.033 | 163.898 | *Rhizoplaca macleanii* (Dodge) Castello | 03 | MK970665 | MN023039 | MN023053 | Tr_A02 | 02 | MK970698 | - | - |
| T46651 | 4 | 374 | -78.028 | 163.851 | *Carbonea vorticosa* (Flörke) Hertel | 01 | MK970656 | - | - | - | - | - | - | - |
| T46659 | 4 | 363 | -78.023 | 163.903 | *Lecidella greenii* Ruprecht & Türk | 01 | MK970671 | - | MN023055 | Tr_A02 | 02 | MK970698 | - | - |
| T46672 | 4 | 397 | -78.024 | 163.898 | *Rhizoplaca macleanii* (Dodge) Castello | 01 | MK970663 | - | - | Tr_A02 | 02 | MK970698 | - | - |
| T46673 | 4 | 365 | -78.027 | 163.851 | *Rhizoplaca macleanii* (Dodge) Castello | 01 | MK970663 | - | - | Tr_A02 | 01 | MK970699 | - | - |
| T46676 | 4 | 359 | -78.028 | 163.851 | *Lecidella greenii* Ruprecht & Türk | 02 | MK970671 | - | - | Tr_A02 | 01 | MK970699 | - | - |
| T46677 | 4 | 843 | -78.027 | 163.848 | *Lecidea cancriformis* Dodge & Baker | 04 | MK970681 | - | - | Tr_A02 | 05 | MK970696 | - | - |
| T46678 | 4 | 843 | -78.028 | 163.850 | *Rhizoplaca macleanii* (Dodge) Castello | 08 | MK970669 | - | - | Tr_A02 | 01 | MK970699 | - | - |
| T46679 | 4 | 790 | -78.036 | 163.971 | *Rhizoplaca macleanii* (Dodge) Castello | 04 | MK970664 | MN023037 | - | Tr_A02 | 05 | MK970696 | - | - |
| T46680 | 4 | 360 | -78.036 | 163.971 | *Lecidea polypycnidophora* Ruprecht & Türk | 01 | MK970663 | MN023043 | MN023053 | Tr_A02 | 01 | MK970699 | - | - |
| T46681 | 4 | 376 | -78.032 | 163.951 | *Rhizoplaca macleanii* (Dodge) Castello | 01 | MK970663 | - | - | Tr_A02 | 02 | MK970698 | - | - |
| T46683 | 4 | 381 | -78.028 | 163.824 | *Huea* sp. 1 | 01 | MK970690 | - | MN023048 | Tr_A02 | 02 | MK970698 | - | - |
| T46684 | 4 | 870 | -78.044 | 163.986 | *Lecidea cancriformis* Dodge & Baker | 01 | MK970677 | - | - | Tr_S02 | 01 | MK970693 | - | - |
| T46685 | 4 | 870 | -78.044 | 163.986 | *Lecidea cancriformis* Dodge & Baker | 01 | MK970677 | - | MN023056 | Tr_S02 | 01 | MK970693 | - | MN023030 |
| T46687 | 4 | 340 | -78.028 | 163.853 | *Huea* sp. 1 | 01 | MK970690 | - | - | Tr_A02 | 02 | MK970698 | - | - |
| T46699 | 4 | 405 | -78.020 | 163.815 | *Huea* sp. 1 | 01 | MK970690 | MN023041 | - | Tr_A02 | 01 | MK970699 | - | - |
| T46701 | 4 | 390 | -78.020 | 163.805 | *Rhizoplaca macleanii* (Dodge) Castello | 01 | MK970663 | MN023034 | - | Tr_A02 | 02 | MK970698 | - | - |
| T46706 | 4 | 379 | -78.028 | 163.821 | *Lecidella greenii* Ruprecht & Türk | 02 | MK970671 | - | MN023054 | Tr_A02 | 02 | MK970698 | - | - |
| T46709 | 3 | 600 | -78.073 | 163.717 | *Buellia* sp. 3 | 01 | MK970688 | MN023047 | - | Tr_A02 | 02 | MK970698 | - | - |
| T46710 | 3 | 600 | -78.073 | 163.717 | *Rhizoplaca macleanii* (Dodge) Castello | 02 | MK970666 | - | - | Tr_A02 | 02 | MK970698 | - | - |
| T46713 | 4 | 350 | -78.028 | 163.843 | *Lecidea polypycnidophora* Ruprecht & Türk | 01 | MK970663 | MN023043 | MN023053 | Tr_A02 | 02 | MK970698 | - | - |
| T46716 | 4 | 412 | -78.040 | 163.802 | *Lecidea polypycnidophora* Ruprecht & Türk | 01 | MK970663 | MN023043 | MN023053 | Tr_A02 | 02 | MK970698 | - | - |
| T46717 | 4 | 442 | -78.040 | 163.806 | *Lecidea polypycnidophora* Ruprecht & Türk | 01 | MK970663 | - | - | Tr_A02 | 02 | MK970698 | - | - |
| T46718 | 4 | 442 | -78.040 | 163.807 | *Carbonea vorticosa* (Flörke) Hertel | 01 | MK970656 | - | - | Tr_A02 | 02 | MK970698 | - | - |
| T46719 | 4 | 425 | -78.038 | 163.804 | *Carbonea* sp. URm1 | 01 | MK970657 | - | - | Tr_A02 | 02 | MK970698 | - | - |
| T46726 | 3 | 890 | -78.071 | 163.785 | *Austrolecia* sp. 4 | 01 | MK970653 | - | - | Tr_A02 | 01 | MK970699 | - | - |
| T48769 | 1 | 370 | -78.126 | 163.700 | *Lecidella greenii* Ruprecht & Türk | 02 | MK970671 | - | - | Tr_A02 | 01 | MK970699 | - | - |
| T48770 | 1 | 373 | -78.127 | 163.690 | *Lecanora* sp. 3 | 01 | MK970659 | - | - | Tr_A02 | 06 | MK970701 | - | - |
| T48772 | 1 | 393 | -78.120 | 163.684 | *Austrolecia* sp. 1 | 01 | MK970650 | - | - | Tr_A02 | 01 | MK970699 | - | - |
| T48773 | 1 | 377 | -78.127 | 163.674 | *Lecidea polypycnidophora* Ruprecht & Türk | 01 | MK970663 | MN023043 | - | Tr_A02 | 01 | MK970699 | MN023065 | - |
| T48774 | 1 | 526 | -78.135 | 163.626 | *Rhizoplaca macleanii* (Dodge) Castello | 01 | MK970663 | - | - | Tr_A02 | 02 | MK970698 | - | - |
| T48776 | 1 | 783 | -78.165 | 163.753 | *Lecidea cancriformis* Dodge & Baker | 02 | MK970679 | MN023046 | MN023057 | Tr_A02 | 03 | MK970702 | - | - |
| T48777 | 1 | 766 | -78.166 | 163.755 | *Lecidea cancriformis* Dodge & Baker | 02 | MK970679 | MN023046 | MN023058 | Tr_S15 | 01 | MK970692 | MN023065 | MN023031 |
| T48778a | 1 | 700 | -78.149 | 163.769 | *Rhizoplaca macleanii* (Dodge) Castello | 01 | MK970663 | - | - | Tr_A02 | 02 | MK970698 | - | - |
| T48778b | 1 | 700 | -78.149 | 163.769 | *Austrolecia* sp. 1 | 01 | MK970650 | - | - | Tr_A02 | 03 | MK970702 | - | - |
| T48779 | 1 | 757 | -78.164 | 163.755 | *Rhizoplaca macleanii* (Dodge) Castello | 04 | MK970664 | MN023037 | - | Tr_A02 | 05 | MK970696 | - | - |
| T48781 | 1 | 745 | -78.128 | 163.620 | *Lecidea* sp. 6 | 02 | MK620097 | - | - | Tr_A02 | 01 | MK970699 | - | - |
| T48782 | 1 | 679 | -78.123 | 163.642 | *Lecidea cancriformis* Dodge & Baker | 02 | MK970679 | MN023046 | - | Tr_S18 | 01 | MK970695 | MN023070 | MN023032 |
| T48783 | 1 | 621 | -78.122 | 163.653 | *Huea* sp. 1 | 01 | MK970690 | MN023041 | MN023048 | Tr_A02 | 01 | MK970699 | - | - |
| T48784 | 1 | 423 | -78.121 | 163.683 | *Lecidea polypycnidophora* Ruprecht & Türk | 01 | MK970663 | MN023043 | - | Tr_A02 | 01 | MK970699 | MN023065 | - |
| T48785 | 1 | 429 | -78.133 | 163.666 | *Lecidella greenii* Ruprecht & Türk | 01 | MK970671 | - | MN023054 | Tr_S15 | 01 | MK970692 | - | - |
| T48786 | 1 | 448 | -78.167 | 163.651 | *Austrolecia* sp. 1 | 01 | MK970650 | - | - | Tr_A02 | 01 | MK970699 | - | - |
| T48787 | 1 | 361 | -78.127 | 163.678 | *Lecidella greenii* Ruprecht & Türk | 01 | MK970671 | - | MN023054 | Tr_A02 | 01 | MK970699 | - | - |
| T48788 | 1 | 397 | -78.120 | 163.684 | *Carbonea vorticosa* (Flörke) Hertel | 01 | MK970656 | MN023033 | MN023050 | Tr_A02 | 01 | MK970699 | - | - |
| T48789 | 1 | 860 | -78.152 | 163.739 | *Carbonea* sp. 2 | 01 | MK970654 | - | MN023051 | Tr_S15 | 01 | MK970692 | MN023069 | MN023031 |
| T48790a | 1 | 415 | -78.120 | 163.682 | *Carbonea vorticosa* (Flörke) Hertel | 01 | MK970656 | MN023033 | - | Tr_A02 | 01 | MK970699 | - | - |
| T48790b | 1 | 415 | -78.120 | 163.682 | *Lecanora* sp. 3 | 01 | MK970659 | - | - | Tr_A02 | 12 | MK970703 | - | - |
| T48791a | 1 | 373 | -78.120 | 163.686 | *Lecidella greenii* Ruprecht & Türk | 01 | MK970671 | - | - | Tr_A02 | 01 | MK970699 | - | - |
| T48792 | 1 | 415 | -78.120 | 163.682 | *Austrolecia* sp. 1 | 01 | MK970650 | - | - | Tr_A02 | 02 | MK970698 | - | - |
| T48793a | 1 | 831 | -78.153 | 163.731 | *Lecidea cancriformis* Dodge & Baker | 01 | MK970677 | - | MN023056 | Tr_S15 | 01 | MK970692 | - | MN023031 |
| T48793b | 1 | 831 | -78.153 | 163.731 | *Carbonea* sp. 2 | 01 | MK970654 | - | - | Tr_S15 | 01 | MK970692 | MN023069 | MN023031 |
| T48793c | 1 | 831 | -78.153 | 163.731 | *Austrolecia* sp. 1 | 01 | MK970650 | - | - | Tr_A02 | 01 | MK970699 | - | - |
| T48794a | 1 | 905 | -78.151 | 163.735 | *Austrolecia* sp. 1 | 01 | MK970650 | - | - | Tr_S15 | 01 | MK970692 | MN023069 | MN023031 |
| T48794b | 1 | 905 | -78.151 | 163.735 | *Rhizoplaca macleanii* (Dodge) Castello | 01 | MK970663 | - | - | Tr_A02 | 02 | MK970698 | - | - |
| T48795a | 1 | 853 | -78.161 | 163.714 | *Rhizoplaca macleanii* (Dodge) Castello | 07 | MK970668 | - | - | Tr_A02 | 02 | MK970698 | - | - |
| T48795b | 1 | 853 | -78.161 | 163.714 | *Austrolecia* sp. 1 | 01 | MK970650 | - | - | Tr_A02 | 02 | MK970698 | - | - |
| T48795c | 1 | 853 | -78.161 | 163.714 | *Austrolecia* sp. 1 | 01 | MK970650 | - | - | Tr_S15 | 01 | MK970692 | MN023069 | - |
| T48796 | 1 | 831 | -78.159 | 163.698 | *Austrolecia* sp. 1 | 01 | MK970650 | - | - | Tr_A02 | 02 | MK970698 | - | - |
| T48797 | 1 | 814 | -78.156 | 163.689 | *Rhizoplaca macleanii* (Dodge) Castello | 05 | MK970667 | - | - | Tr_A02 | 02 | MK970698 | - | - |
| T48798 | 1 | 911 | -78.150 | 163.736 | *Rhizoplaca macleanii* (Dodge) Castello | 05 | MK970667 | - | - | Tr_A02 | 02 | MK970698 | - | - |
| T48799b | 1 | 743 | -78.145 | 163.620 | *Lecidea cancriformis* Dodge & Baker | 02 | MK970679 | - | - | Tr_A02 | 02 | MK970698 | - | - |
| T48799c | 1 | 743 | -78.145 | 163.620 | *Lecidea cancriformis* Dodge & Baker | 01 | MK970677 | - | - | Tr_S15 | 01 | MK970692 | - | - |
| T48800 | 1 | 729 | -78.146 | 163.631 | *Lecanora* sp. 2 | 01 | MK970662 | MN023036 | MN023052 | Tr_A02 | 02 | MK970698 | - | - |
| T48801a | 1 | 732 | -78.144 | 163.626 | *Lecanora* sp. 3 | 01 | MK970659 | - | - | Tr_A02 | 01 | MK970699 | - | - |
| T48801c | 1 | 732 | -78.144 | 163.626 | *Lecidella greenii* Ruprecht & Türk | 01 | MK970671 | MN023040 | - | Tr_A02 | 01 | MK970699 | - | - |
| T48802 | 1 | 732 | -78.147 | 163.629 | *Austrolecia* sp. 1 | 01 | MK970650 | - | - | Tr_A02 | 01 | MK970699 | - | - |
| T48803a | 1 | 775 | -78.148 | 163.630 | *Austrolecia* sp. 1 | 01 | MK970650 | - | - | Tr_A02 | 01 | MK970699 | - | - |
| T48803b | 1 | 775 | -78.148 | 163.630 | *Carbonea* sp. URm1 | 01 | MK970657 | MN023034 | - | Tr_A02 | 01 | MK970699 | - | - |
| T48804 | 1 | 775 | -78.148 | 163.630 | *Rhizoplaca macleanii* (Dodge) Castello | 09 | MK970670 | - | MN023053 | Tr_A02 | 01 | MK970699 | - | - |
| T48805 | 1 | 732 | -78.144 | 163.626 | *Lecidella greenii* Ruprecht & Türk | 01 | MK970671 | - | - | Tr_A02 | 02 | MK970698 | - | - |
| T48806 | 1 | 568 | -78.142 | 163.628 | *Carbonea* sp. URm1 | 01 | MK970657 | MN023034 | - | Tr_A02 | 01 | MK970699 | - | - |
| T48807 | 1 | 554 | -78.141 | 163.631 | *Lecidea andersonii* Filson | 01 | MK970673 | MN023042 | MN023060 | Tr_A02 | 03 | MK970702 | - | - |
| T48809 | 1 | 540 | -78.148 | 163.657 | *Lecidella greenii* Ruprecht & Türk | 01 | MK970671 | - | - | Tr_A02 | 01 | MK970699 | - | - |
| T48811a | 1 | 557 | -78.142 | 163.657 | *Lecidella greenii* Ruprecht & Türk | 01 | MK970671 | - | MN023055 | Tr_A02 | 02 | MK970698 | - | - |
| T48812a | 1 | 540 | -78.138 | 163.619 | *Lecidella greenii* Ruprecht & Türk | 01 | MK970671 | - | - | Tr_A02 | 01 | MK970699 | - | - |
| T48812b | 1 | 540 | -78.138 | 163.619 | *Lecidella greenii* Ruprecht & Türk | 01 | MK970671 | - | MN023055 | Tr_A02 | 01 | MK970699 | - | - |
| T48813 | 1 | 557 | -78.142 | 163.657 | *Lecidea* UCR1 | 01 | MK970675 | MN023044 | - | Tr_A02 | 01 | MK970699 | - | - |
| T48815 | 1 | 516 | -78.137 | 163.627 | *Austrolecia* sp. 1 | 02 | MK970652 | - | - | Tr_A02 | 02 | MK970698 | - | - |
| T48816 | 2 | 171 | -78.100 | 163.810 | *Buellia* sp. 1 | 01 | MK970689 | - | - | Tr_A02 | 02 | MK970698 | - | - |
| T48817a | 2 | 546 | -78.113 | 163.785 | *Rhizoplaca macleanii* (Dodge) Castello | 01 | MK970663 | - | - | Tr_A02 | 02 | MK970698 | - | - |
| T48817b | 2 | 546 | -78.113 | 163.785 | *Carbonea* sp. 2 | 02 | MK970655 | - | - | Tr_A02 | 03 | MK970702 | - | - |
| T48818 | 1 | 596 | -78.148 | 163.811 | *Austrolecia* sp. 1 | 03 | MK970651 | - | - | Tr_A02 | 01 | MK970699 | - | - |
| T48820 | 2 | 575 | -78.114 | 163.780 | *Carbonea* sp. 2 | 01 | MK970654 | - | - | Tr_A02 | 02 | MK970698 | - | - |
| T48821 | 2 | 589 | -78.114 | 163.779 | *Rhizoplaca macleanii* (Dodge) Castello | 03 | MK970665 | - | - | Tr_A02 | 02 | MK970698 | - | - |
| T48823a | 2 | 401 | -78.098 | 163.710 | *Carbonea vorticosa* (Flörke) Hertel | 01 | MK970656 | - | - | - | - | - | - | - |
| T48824 | 2 | 345 | -78.097 | 163.707 | *Buellia* sp. 1 | 01 | MK970689 | - | - | Tr_A02 | 01 | MK970699 | - | - |
| T48825 | 2 | 498 | -78.097 | 163.691 | *Carbonea* sp. 2 | 01 | MK970654 | MN023035 | - | Tr_A02 | 01 | MK970699 | - | - |
| T48826 | 2 | 424 | -78.097 | 163.717 | *Lecanora fuscobrunnea* Dodge & Baker | 01 | MK970661 | - | - | Tr_S02 | 02 | MK970694 | MN023068 | - |
| T48827a | 2 | 345 | -78.097 | 163.707 | *Buellia frigida* Darbishire | 04 | MK970687 | - | - | Tr_A02 | 01 | MK970699 | - | - |
| T48827b | 2 | 345 | -78.097 | 163.707 | *Huea* sp. 1 | 01 | MK970690 | MN023041 | - | Tr_A02 | 03 | MK970702 | - | - |
| T48828a | 2 | 406 | -78.110 | 163.858 | *Lecidella greenii* Ruprecht & Türk | 02 | MK970671 | - | MN023054 | - | - | - | - | - |
| T48828b | 2 | 406 | -78.110 | 163.858 | *Lecidella greenii* Ruprecht & Türk | 01 | MK970671 | - | - | Tr_A02 | 02 | MK970698 | - | - |
| T48829 | 2 | 453 | -78.111 | 163.858 | *Lecidella greenii* Ruprecht & Türk | 01 | MK970671 | - | - | Tr_A02 | 01 | MK970699 | - | - |
| T48831 | 2 | 451 | -78.111 | 163.858 | *Carbonea vorticosa* (Flörke) Hertel | 01 | MK970656 | MN023033 | - | Tr_A02 | 01 | MK970699 | - | - |
| T48832 | 2 | 526 | -78.112 | 163.824 | *Carbonea* sp. 2 | 01 | MK970654 | - | MN023051 | Tr_A02 | 03 | MK970702 | - | - |
| T48834 | 2 | 513 | -78.114 | 163.854 | *Lecidea cancriformis* Dodge & Baker | 01 | MK970677 | - | - | Tr_S15 | 01 | MK970692 | - | - |
| T48835 | 2 | 522 | -78.112 | 163.820 | *Buellia* sp. 1 | 01 | MK970689 | - | - | Tr_A02 | 01 | MK970699 | - | - |
| T48836 | 2 | 179 | -78.099 | 163.778 | *Lecanora cf. mons-nivis* Darbishire | 01 | MK970658 | - | - | Tr_A02 | 01 | MK970699 | - | - |
| T48837 | 2 | 179 | -78.098 | 163.777 | *Lecidella greenii* Ruprecht & Türk | 01 | MK970671 | - | MN023055 | Tr_A02 | 01 | MK970699 | - | - |
| T48839 | 2 | 514 | -78.114 | 163.854 | *Lecidella greenii* Ruprecht & Türk | 01 | MK970671 | - | - | Tr_A02 | 01 | MK970699 | - | - |
| T48840 | 3 | 722 | -78.068 | 163.861 | *Buellia* sp. 1 | 01 | MK970689 | - | - | Tr_A02 | 02 | MK970698 | - | - |
| T48841a | 3 | 722 | -78.068 | 163.861 | *Rhizoplaca macleanii* (Dodge) Castello | 01 | MK970663 | - | - | Tr_A02 | 02 | MK970698 | - | - |
| T48841b | 3 | 722 | -78.068 | 163.861 | *Carbonea vorticosa* (Flörke) Hertel | 01 | MK970656 | - | MN023050 | Tr_A02 | 02 | MK970698 | - | - |
| T48843a | 3 | 836 | -78.066 | 163.870 | *Lecidea cancriformis* Dodge & Baker | 01 | MK970677 | - | - | Tr_S02 | 01 | MK970693 | - | - |
| T48843b | 3 | 836 | -78.066 | 163.870 | *Austrolecia* sp. 1 | 01 | MK970650 | - | - | Tr_S15 | 01 | MK970692 | MN023069 | - |
| T48843c | 3 | 836 | -78.066 | 163.870 | *Lecidea cancriformis* Dodge & Baker | 05 | MK970678 | - | - | - | - | - | - | - |
| T48843d | 3 | 836 | -78.066 | 163.870 | *Rhizoplaca macleanii* (Dodge) Castello | 01 | MK970663 | - | - | Tr_A02 | 01 | MK970699 | - | - |
| T48843e | 3 | 836 | -78.066 | 163.870 | *Rhizoplaca macleanii* (Dodge) Castello | 01 | MK970663 | - | - | Tr_A02 | 03 | MK970702 | - | - |
| T48844 | 3 | 749 | -78.067 | 163.863 | *Lecidella greenii* Ruprecht & Türk | 02 | MK970671 | - | - | Tr_A02 | 02 | MK970698 | - | - |
| T48849 | 3 | 509 | -78.077 | 163.783 | *Austrolecia* sp. 1 | 01 | MK970650 | - | - | Tr_A02 | 01 | MK970699 | - | - |
| T48851 | 3 | 587 | -78.074 | 163.793 | *Lecanora fuscobrunnea* Dodge & Baker | 02 | MK970660 | MN023037 | - | Tr_A02 | 02 | MK970698 | - | - |
| T48852 | 3 | 587 | -78.075 | 163.792 | *Huea* sp. 1 | 01 | MK970690 | - | - | Tr_A02 | 01 | MK970699 | - | - |
| T48853 | 3 | 594 | -78.075 | 163.791 | *Huea* sp. 1 | 01 | MK970690 | - | MN023048 | Tr_A02 | 01 | MK970699 | - | - |
| T48855a | 4 | 688 | -78.036 | 163.837 | *Austrolecia* sp. 1 | 01 | MK970650 | - | - | Tr_A02 | 01 | MK970699 | - | - |
| T48855b | 4 | 688 | -78.036 | 163.837 | *Lecidea cancriformis* Dodge & Baker | 03 | MK970680 | MN023046 | MN023059 | Tr_S15 | 01 | MK970692 | MN023069 | MN023031 |
| T48857a | 4 | 688 | -78.036 | 163.827 | *Lecidea* sp. 6 | 01 | MK970684 | MN023045 | - | Tr_A02 | 01 | MK970699 | - | - |
| T48857b | 4 | 688 | -78.036 | 163.827 | *Buellia* sp. 1 | 01 | MK970689 | - | - | Tr_S15 | 01 | MK970692 | - | MN023031 |
| T48857c | 4 | 688 | -78.036 | 163.827 | *Austrolecia* sp. 1 | 01 | MK970650 | - | - | Tr_A02 | 02 | MK970698 | - | - |
| T48858 | 4 | 319 | -78.025 | 163.975 | *Lecidella greenii* Ruprecht & Türk | 01 | MK970671 | - | - | - | - | - | - | - |
| T48859 | 4 | 319 | -78.025 | 163.986 | *Lecidea polypycnidophora* Ruprecht & Türk | 01 | MK970663 | - | MN023053 | Tr_A02 | 02 | MK970698 | - | - |
| T48860a | 4 | 692 | -78.036 | 163.836 | *Huea* sp. 1 | 01 | MK970690 | - | MN023048 | Tr_A02 | 01 | MK970699 | - | - |
| T48860b | 4 | 694 | -78.036 | 163.836 | *Lecidea* sp. 6 | 01 | MK970684 | MN023045 | MN023064 | Tr_A02 | 01 | MK970699 | - | - |
| T48860c | 4 | 694 | -78.036 | 163.836 | *Huea* sp. 1 | 01 | MK970690 | - | - | Tr_A02 | 02 | MK970698 | - | - |
| T48860d | 4 | 694 | -78.036 | 163.836 | *Huea* sp. 1 | 01 | MK970690 | - | - | Tr_A02 | 01 | MK970699 | - | - |
| T48861 | 4 | 380 | -78.024 | 163.892 | *Lecanora* sp. 3 | 01 | MK970659 | - | - | Tr_A02 | 01 | MK970699 | - | - |
| T48862 | 4 | 875 | -78.037 | 163.978 | *Lecidea cancriformis* Dodge & Baker | 06 | MK970682 | - | - | - | - | - | - | - |
| T48864 | 4 | 850 | -78.039 | 163.989 | *Lecidea cancriformis* Dodge & Baker | 02 | MK970679 | - | - | Tr_A02 | 03 | MK970702 | - | - |
| T48865a | 4 | 874 | -78.036 | 163.990 | *Rhizoplaca macleanii* (Dodge) Castello | 02 | MK970666 | - | - | Tr_A02 | 02 | MK970698 | MN023067 | - |
| T48867a | 4 | 936 | -78.043 | 164.104 | *Rhizoplaca macleanii* (Dodge) Castello | 01 | MK970663 | - | - | Tr_A02 | 05 | MK970696 | - | - |
| T48867b | 4 | 936 | -78.043 | 164.104 | *Austrolecia* sp. 1 | 01 | MK970650 | - | - | Tr_A02 | 01 | MK970699 | - | - |
| T48867c | 4 | 936 | -78.043 | 164.104 | *Rhizoplaca macleanii* (Dodge) Castello | 02 | MK970666 | - | - | Tr_A02 | 03 | MK970702 | - | - |
| T48869 | 4 | 849 | -78.037 | 163.978 | *Lecidella greenii* Ruprecht & Türk | 01 | MK970671 | - | - | - | - | - | - | - |
| T48872 | 4 | 673 | -78.030 | 163.951 | *Lecidella greenii* Ruprecht & Türk | 01 | MK970671 | - | - | - | - | - | - | - |
| T48873 | 4 | 455 | -78.024 | 163.893 | *Lecidella greenii* Ruprecht & Türk | 01 | MK970671 | - | MN023054 | Tr_A02 | 02 | MK970698 | - | - |
| T48874 | 4 | 375 | -78.024 | 163.899 | *Lecidea* UCR1 | 02 | MK970676 | MN023044 | MN023062 | Tr_A02 | 11 | MK970697 | MN023066 | - |
| T48875 | 4 | 375 | -78.024 | 163.900 | *Lecidea lapicida* (Ach.) Ach. subsp. | 01 | MK970683 | - | - | Tr_A02 | 01 | MK970699 | - | - |
| T48876a | 4 | 380 | -78.024 | 163.900 | *Lecidella greenii* Ruprecht & Türk | 01 | MK970671 | MN023040 | - | Tr_A02 | 02 | MK970698 | - | - |
| T48876b | 4 | 380 | -78.024 | 163.900 | *Lecidella greenii* Ruprecht & Türk | 01 | MK970671 | - | MN023055 | Tr_A02 | 01 | MK970699 | - | - |
| T48877 | 4 | 385 | -78.024 | 163.900 | *Carbonea vorticosa* (Flörke) Hertel | 01 | MK970656 | - | MN023050 | Tr_A02 | 02 | MK970698 | - | - |
| T48879 | 3 | 419 | -78.057 | 163.747 | *Lecidella greenii* Ruprecht & Türk | 01 | MK970671 | - | MN023055 | - | - | - | - | - |
| T48880 | 3 | 443 | -78.058 | 163.740 | *Lecidea polypycnidophora* Ruprecht & Türk | 01 | MK970674 | - | - | - | - | - | - | - |
| T48881a | 3 | 541 | -78.061 | 163.791 | *Huea* sp. 1 | 01 | MK970690 | - | MN023049 | Tr_A02 | 03 | MK970702 | - | - |
| T48881b | 3 | 541 | -78.061 | 163.791 | *Rhizoplaca macleanii* (Dodge) Castello | 02 | MK970666 | - | - | Tr_A02 | 01 | MK970699 | - | - |
| T48882 | 3 | 575 | -78.057 | 163.817 | *Lecidea polypycnidophora* Ruprecht & Türk | 01 | MK970663 | - | - | Tr_A02 | 01 | MK970699 | MN023065 | - |
| T48883a | 3 | 671 | -78.058 | 163.847 | *Huea* sp. 1 | 01 | MK970690 | - | - | Tr_A02 | 02 | MK970698 | - | - |
| T48883b | 3 | 671 | -78.058 | 163.847 | *Lecidea* sp. 5 | 01 | MK620099 | - | MN023063 | Tr_A02 | 09 | MK970700 | - | - |
| T48885 | 3 | 666 | -78.057 | 163.844 | *Rhizoplaca macleanii* (Dodge) Castello | 02 | MK970666 | - | - | Tr_A02 | 02 | MK970698 | - | - |
| T48886 | 4 | 464 | -78.025 | 163.932 | *Austrolecia* sp. 1 | 01 | MK970650 | - | - | Tr_A02 | 02 | MK970698 | - | - |
| T48887a | 3 | 710 | -78.070 | 163.711 | *Lecidea cancriformis* Dodge & Baker | 02 | MK970679 | MN023046 | - | Tr_S15 | 01 | MK970692 | MN023069 | - |
| T48887b | 3 | 710 | -78.070 | 163.711 | *Huea* sp. 1 | 01 | MK970690 | - | - | Tr_A02 | 03 | MK970702 | - | - |
| T48888b | 3 | 629 | -78.073 | 163.717 | *Lecanora cf. mons-nivis* Darbishire | 01 | MK970658 | - | - | Tr_A02 | 01 | MK970699 | - | - |
| T48900 | 4 | 610 | -78.034 | 163.845 | *Rhizoplaca macleanii* (Dodge) Castello | 01 | MK970663 | - | - | Tr_A02 | 02 | MK970698 | - | - |

**Online Resource 1b**: Additional samples taken from Perez-Ortega et al. (2012) and used in this study, with information on haplotypes, collecting localities and Genbank accession numbers.

| Laboratory code | area | elevation | latitude | longitude | Mycobiont |  |  | Associated green micro algae |  |  |
| --- | --- | --- | --- | --- | --- | --- | --- | --- | --- | --- |
|  |  |  |  |  | species_name | HT | Accession numbers nrITS | OTU_id | HT | Accession numbers nrITS |
| s106 | 2 | 566 | -78.113 | 163.782 | *Lecanora* sp. 2 | 01 | JX036037 | Tr_A02 | 02 | JX036159 |
| s111 | 2 | 520 | -78.114 | 163.854 | *Buellia frigida* Darbshire | 01 | JX036042 | Tr_A02 | 01 | JX036164 |
| s112 | 2 | 520 | -78.114 | 163.854 | *Buellia* sp. 2 | 01 | JX036043 | Tr_A02 | 01 | JX036165 |
| s113 | 2 | 520 | -78.114 | 163.854 | *Lecidea cancriformis* Dodge & Baker | 01 | JX036044 | Tr_S15 | 01 | JX036166 |
| s114 | 2 | 520 | -78.114 | 163.854 | *Lecidea cancriformis* Dodge & Baker | 01 | JX036045 | Tr_A02 | 05 | JX036167 |
| s115 | 3 | 839 | -78.066 | 163.870 | *Lecidella greenii* Ruprecht & Türk | 02 | JX036046 | Tr_A02 | 03 | JX036168 |
| s117 | 3 | 839 | -78.066 | 163.870 | *Buellia frigida* Darbshire | 01 | JX036048 | Tr_A02 | 03 | JX036169 |
| s118 | 3 | 839 | -78.066 | 163.870 | *Buellia frigida* Darbshire | 01 | JX036049 | Tr_A02 | 02 | JX036170 |
| s119 | 3 | 411 | -78.083 | 163.768 | *Austrolecia* sp. 1 | 01 | JX036050 | Tr_A02 | 01 | JX036171 |
| s120 | 3 | 411 | -78.083 | 163.768 | *Lecidea polypycnidophora* Ruprecht & Türk | 01 | JX036051 | Tr_A02 | 02 | JX036172 |
| s121 | 4 | 384 | -78.024 | 163.900 | *Carbonea vorticosa* (Flörke) Hertel | 01 | JX036052 | Tr_A02 | 01 | JX036173 |
| s122 | 2 | 413 | -78.110 | 163.787 | *Carbonea vorticosa* (Flörke) Hertel | 01 | JX036053 | Tr_A02 | 01 | JX036174 |
| s123 | 2 | 413 | -78.110 | 163.787 | *Carbonea vorticosa* (Flörke) Hertel | 01 | JX036054 | Tr_A02 | 01 | JX036175 |
| s124 | 3 | 666 | -78.057 | 163.844 | *Rhizoplaca macleanii* (Dodge) Castello | 02 | JX036055 | Tr_A02 | 02 | JX036176 |
| s125 | 3 | 666 | -78.057 | 163.844 | *Rhizoplaca macleanii* (Dodge) Castello | 02 | JX036056 | Tr_A02 | 02 | JX036177 |
| s126 | 3 | 541 | -78.061 | 163.791 | *Huea* sp. 1 | 01 | JX036057 | Tr_A02 | 03 | JX036178 |
| s127 | 4 | 874 | -78.036 | 163.990 | *Buellia frigida* Darbshire | 01 | JX036058 | Tr_A02 | 07 | JX036179 |
| s128 | 3 | 581 | -78.075 | 163.790 | *Austrolecia* sp. 1 | 01 | JX036059 | Tr_A02 | 10 | JX036180 |
| s165 | 2 | 465 | -78.097 | 163.691 | *Buellia frigida* Darbshire | 01 | JX036070 | Tr_A02 | 02 | JX036191 |
| s166 | 2 | 597 | -78.097 | 163.691 | *Buellia frigida* Darbshire | 01 | JX036071 | Tr_A02 | 01 | JX036192 |
| s168 | 4 | 688 | -78.036 | 163.837 | *Austrolecia* sp. 1 | 02 | JX036073 | Tr_A02 | 04 | JX036194 |
| s171 | 4 | 688 | -78.036 | 163.837 | *Lecanora* sp. 2 | 01 | JX036076 | Tr_A02 | 01 | JX036197 |
| s173 | 2 | 521 | -78.114 | 163.854 | *Lecanora* sp. 2 | 01 | JX036078 | Tr_A02 | 01 | JX036199 |
| s174 | 4 | 597 | -78.033 | 163.849 | *Buellia frigida* Darbshire | 01 | JX036079 | Tr_A02 | 02 | JX036200 |
| s175 | 4 | 597 | -78.033 | 163.849 | *Lecanora* sp. 2 | 01 | JX036080 | Tr_A02 | 04 | JX036201 |
| s176 | 3 | 658 | -78.073 | 163.712 | *Buellia frigida* Darbshire | 01 | JX036081 | Tr_A02 | 01 | JX036202 |
| s177 | 2 | 521 | -78.114 | 163.854 | *Austrolecia* sp. 2 | 01 | JX036082 | Tr_A02 | 01 | JX036203 |
| s178 | 2 | 521 | -78.114 | 163.854 | *Austrolecia* sp. 2 | 02 | JX036083 | Tr_A02 | 01 | JX036204 |
| s179 | 3 | 540 | -78.063 | 163.809 | *Lecanora* sp. 2 | 01 | JX036084 | Tr_A02 | 01 | JX036205 |
| s180 | 3 | 540 | -78.063 | 163.809 | *Buellia frigida* Darbshire | 01 | JX036085 | Tr_A02 | 01 | JX036206 |
| s181 | 2 | 453 | -78.111 | 163.858 | *Lecidella greenii* Ruprecht & Türk | 01 | JX036086 | Tr_A02 | 04 | JX036207 |
| s182 | 4 | 688 | -78.036 | 163.837 | *Austrolecia* sp. 1 | 01 | JX036087 | Tr_A02 | 06 | JX036208 |
| s184 | 3 | 615 | -78.073 | 163.717 | *Austrolecia* sp. 1 | 01 | JX036089 | Tr_A02 | 02 | JX036210 |
| s187 | 2 | 575 | -78.114 | 163.778 | *Buellia frigida* Darbshire | 02 | JX036092 | Tr_A02 | 13 | JX036213 |
| s189 | 3 | 719 | -78.071 | 163.712 | *Buellia frigida* Darbshire | 01 | JX036094 | Tr_A02 | 02 | JX036215 |
| s190 | 4 | 464 | -78.025 | 163.899 | *Lecanora* sp. 3 | 02 | JX036095 | Tr_A02 | 02 | JX036216 |
| s191 | 4 | 419 | -78.025 | 163.900 | *Lecanora* sp. 3 | 02 | JX036096 | Tr_A02 | 01 | JX036217 |
| s192 | 4 | 481 | -78.030 | 163.834 | *Lecidella greenii* Ruprecht & Türk | 01 | JX036097 | Tr_A02 | 01 | JX036218 |
| s195 | 4 | 874 | -78.036 | 163.992 | *Buellia frigida* Darbshire | 01 | JX036100 | Tr_A02 | 01 | JX036221 |
| s197 | 3 | 541 | -78.061 | 163.791 | *Rhizoplaca macleanii* (Dodge) Castello | 02 | JX036101 | Tr_A02 | 01 | JX036222 |
| s198 | 3 | 541 | -78.061 | 163.791 | *Rhizoplaca macleanii* (Dodge) Castello | 02 | JX036102 | Tr_A02 | 01 | JX036223 |
| s199 | 4 | 375 | -78.024 | 163.900 | *Austrolecia* sp. 1 | 01 | JX036103 | Tr_A02 | 02 | JX036224 |
| s200 | 4 | 375 | -78.024 | 163.900 | *Austrolecia* sp. 1 | 01 | JX036104 | Tr_A02 | 02 | JX036225 |
| s201 | 4 | 375 | -78.024 | 163.900 | *Rhizoplaca macleanii* (Dodge) Castello | 03 | JX036105 | Tr_A02 | 02 | JX036226 |
| s202 | 4 | 360 | -78.027 | 163.839 | *Lecanora* cf. *mons-nivis* Darbshire | 01 | JX036106 | Tr_A02 | 08 | JX036227 |
| s203 | 4 | 360 | -78.027 | 163.839 | *Lecidella greenii* Ruprecht & Türk | 01 | JX036107 | Tr_A02 | 01 | JX036228 |
| s205 | 3 | 722 | -78.068 | 163.861 | *Rhizoplaca macleanii* (Dodge) Castello | 03 | JX036108 | Tr_A02 | 04 | JX036229 |
| s206 | 3 | 722 | -78.068 | 163.861 | *Rhizoplaca macleanii* (Dodge) Castello | 03 | JX036109 | Tr_A02 | 02 | JX036230 |
| s207 | 3 | 722 | -78.068 | 163.861 | *Rhizoplaca macleanii* (Dodge) Castello | 03 | JX036110 | Tr_A02 | 08 | JX036231 |
| s208 | 3 | 722 | -78.068 | 163.861 | *Rhizoplaca macleanii* (Dodge) Castello | 01 | JX036111 | Tr_A02 | 02 | JX036232 |
| s209 | 3 | 722 | -78.068 | 163.861 | *Rhizoplaca macleanii* (Dodge) Castello | 01 | JX036112 | Tr_A02 | 01 | JX036233 |
| s212 | 4 | 710 | -78.034 | 163.845 | *Rhizoplaca macleanii* (Dodge) Castello | 01 | JX036115 | Tr_A02 | 02 | JX036236 |
| s213 | 4 | 710 | -78.034 | 163.845 | *Rhizoplaca macleanii* (Dodge) Castello | 01 | JX036116 | Tr_A02 | 02 | JX036237 |
| s214 | 4 | 710 | -78.034 | 163.845 | *Rhizoplaca macleanii* (Dodge) Castello | 01 | JX036117 | Tr_A02 | 02 | JX036238 |
| s215 | 3 | 710 | -78.070 | 163.711 | *Lecanora fuscobrunnea* Dodge & Baker | 01 | JX036118 | Tr_A02 | 02 | JX036239 |
| s221 | 2 | 508 | -78.112 | 163.823 | *Buellia frigida* Darbshire | 01 | JX036124 | Tr_A02 | 01 | JX036245 |
| s229 | 4 | 710 | -78.034 | 163.845 | *Huea* sp. 1 | 01 | JX036132 | Tr_A02 | 03 | JX036251 |
| s230 | 4 | 710 | -78.034 | 163.845 | *Lecidella greenii* Ruprecht & Türk | 01 | JX036133 | Tr_A02 | 02 | JX036252 |
| s231 | 4 | 710 | -78.034 | 163.845 | *Austrolecia* sp. 1 | 01 | JX036134 | Tr_A02 | 03 | JX036253 |
| s232 | 4 | 710 | -78.034 | 163.845 | *Rhizoplaca macleanii* (Dodge) Castello | 01 | JX036135 | Tr_A02 | 01 | JX036254 |
| s233 | 4 | 710 | -78.034 | 163.845 | *Rhizoplaca macleanii* (Dodge) Castello | 02 | JX036136 | Tr_A02 | 04 | JX036255 |
| s235 | 2 | 589 | -78.113 | 163.778 | *Rhizoplaca macleanii* (Dodge) Castello | 01 | JX036138 | Tr_A02 | 02 | JX036257 |
| s236 | 2 | 589 | -78.113 | 163.778 | *Rhizoplaca macleanii* (Dodge) Castello | 01 | JX036139 | Tr_A02 | 02 | JX036258 |
| s237 | 2 | 589 | -78.113 | 163.778 | *Rhizoplaca macleanii* (Dodge) Castello | 01 | JX036140 | Tr_A02 | 07 | JX036259 |
| s266 | 3 | 583 | -78.075 | 163.791 | *Lecidella greenii* Ruprecht & Türk | 01 | JX036141 | Tr_A02 | 04 | JX036260 |
| s267 | 4 | 597 | -78.033 | 163.849 | *Buellia frigida* Darbshire | 01 | JX036142 | Tr_A02 | 01 | JX036261 |
| s269 | 2 | 566 | -78.113 | 163.782 | *Huea* sp. 1 | 01 | JX036143 | Tr_A02 | 01 | JX036262 |
| s270 | 2 | 566 | -78.113 | 163.782 | *Huea* sp. 1 | 01 | JX036144 | Tr_A02 | 01 | JX036263 |
| s271 | 4 | 959 | -78.047 | 164.104 | *Rhizoplaca macleanii* (Dodge) Castello | 03 | JX036145 | Tr_A02 | 02 | JX036264 |
| s272 | 4 | 809 | -78.035 | 163.978 | *Rhizoplaca macleanii* (Dodge) Castello | 01 | JX036146 | Tr_A02 | 01 | JX036265 |
| s273 | 4 | 688 | -78.030 | 163.949 | *Rhizoplaca macleanii* (Dodge) Castello | 01 | JX036147 | Tr_A02 | 01 | JX036266 |
| s274 | 4 | 874 | -78.036 | 163.990 | *Rhizoplaca macleanii* (Dodge) Castello | 01 | JX036148 | Tr_A02 | 01 | JX036267 |
| s275 | 4 | 874 | -78.036 | 163.990 | *Lecanora* sp. 1 | 01 | JX036149 | Tr_A02 | 03 | JX036268 |
| s300 | 4 | 481 | -78.030 | 163.834 | *Lecidella greenii* Ruprecht & Türk | 01 | JX036150 | Tr_A02 | 01 | JX036269 |
| s301 | 4 | 481 | -78.030 | 163.834 | *Lecidella greenii* Ruprecht & Türk | 01 | JX036151 | Tr_A02 | 01 | JX036270 |
| s95 | 4 | 375 | -78.024 | 163.900 | *Rhizoplaca macleanii* (Dodge) Castello | 06 | JX036152 | Tr_A02 | 04 | JX036271 |
| s96 | 3 | 585 | -78.075 | 163.791 | *Buellia frigida* Darbshire | 03 | JX036153 | Tr_A02 | 05 | JX036272 |
| s97 | 2 | 575 | -78.114 | 163.814 | *Austrolecia* sp. 1 | 01 | JX036154 | Tr_S18 | 01 | JX036273 |

**Online Resource 1c**: The pairs of *min10MycoSp* species and *min10PhoHap* haplotypes that showed significant differences with regards to elevation of sampling sites, as well as the associated p-values of the mctp-tests. The pairs are listed line-by-line; the species with the samples found in higher elevations is given on the left, respectively. PH = photobiont haplotypes

| Elevation of Sample Sites in Pairwise Comparison: |  |  | p-value |
| --- | --- | --- | --- |
|  | higher | lower |  |
| Mycobiont Species | *Austrolecia* sp. 1 | *Carbonea vorticosa* | 0.012 |
|  | *Austrolecia* sp. 1 | *Lecidea polypycnidophora* | 0.001 |
|  | *Buellia frigida* | *Carbonea vorticosa* | 0.027 |
|  | *Buellia frigida* | *Lecidea polypycnidophora* | 0.003 |
|  | *Huea* sp. 1 | *Lecidea polypycnidophora* | 0.049 |
|  | *Lecidea cancriformis* | *Carbonea vorticosa* | 0.000 |
|  | *Lecidea cancriformis* | *Huea* sp. 1 | 0.017 |
|  | *Lecidea cancriformis* | *Lecidea polypycnidophora* | 0.000 |
|  | *Lecidea cancriformis* | *Lecidella greenii* | 0.000 |
|  | *Rhizoplaca macleanii* | *Carbonea vorticosa* | 0.001 |
|  | *Rhizoplaca macleanii* | *Lecidea polypycnidophora* | 0.000 |
|  | *Rhizoplaca macleanii* | *Lecidella greenii* | 0.002 |
| PH | *Tr_A02_h03* | *Tr_A02_h01* | 0.021 |
|  | *Tr_S15_h01* | *Tr_A02_h01* | 0.011 |

**Online Resource 1d** Network matrix giving the number of associations between the mycobiont species and photobiont haplotypes

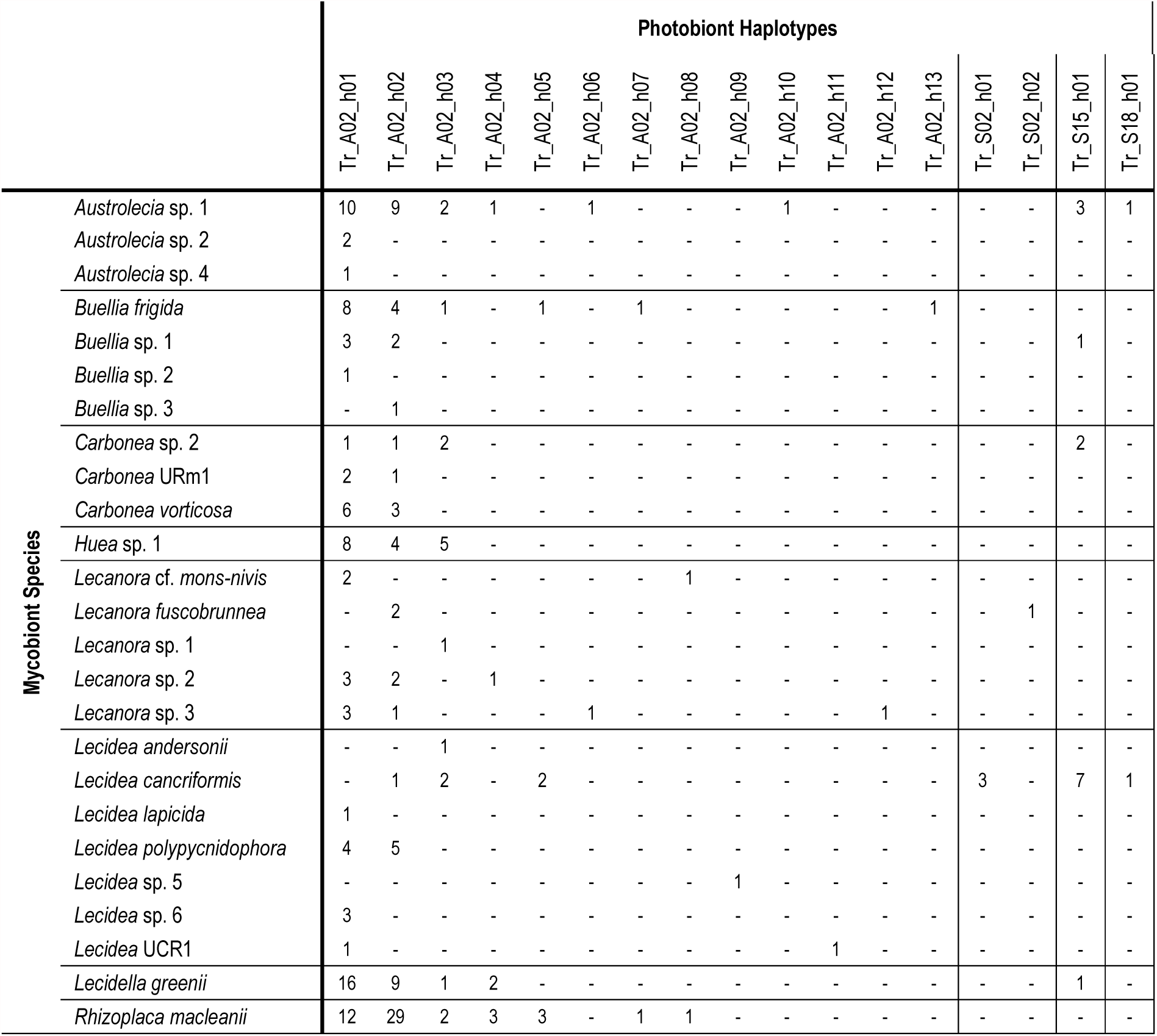

|  |  | Photobiont Haplotypes |  |  |  |  |  |  |  |  |  |  |  |  |  |  |  |  |
| --- | --- | --- | --- | --- | --- | --- | --- | --- | --- | --- | --- | --- | --- | --- | --- | --- | --- | --- |
|  |  | Tr_A02_h01 | Tr_A02_h02 | Tr_A02_h03 | Tr_A02_h04 | Tr_A02_h05 | Tr_A02_h06 | Tr_A02_h07 | Tr_A02_h08 | Tr_A02_h09 | Tr_A02_h10 | Tr_A02_h11 | Tr_A02_h12 | Tr_A02_h13 | Tr_S02_h01 | Tr_S02_h02 | Tr_S15_h01 | Tr_S18_h01 |
| Mycobiont Species | *Austrolecia* sp. 1 | 10 | 9 | 2 | 1 | - | 1 | - | - | - | 1 | - | - | - | - | - | 3 | 1 |
|  | *Austrolecia* sp. 2 | 2 | - | - | - | - | - | - | - | - | - | - | - | - | - | - | - | - |
|  | *Austrolecia* sp. 4 | 1 | - | - | - | - | - | - | - | - | - | - | - | - | - | - | - | - |
|  | *Buellia frigida* | 8 | 4 | 1 | - | 1 | - | 1 | - | - | - | - | - | 1 | - | - | - | - |
|  | *Buellia* sp. 1 | 3 | 2 | - | - | - | - | - | - | - | - | - | - | - | - | - | 1 | - |
|  | *Buellia* sp. 2 | 1 | - | - | - | - | - | - | - | - | - | - | - | - | - | - | - | - |
|  | *Buellia* sp. 3 | - | 1 | - | - | - | - | - | - | - | - | - | - | - | - | - | - | - |
|  | *Carbonea* sp. 2 | 1 | 1 | 2 | - | - | - | - | - | - | - | - | - | - | - | - | 2 | - |
|  | *Carbonea* URm1 | 2 | 1 | - | - | - | - | - | - | - | - | - | - | - | - | - | - | - |
|  | *Carbonea vorticosa* | 6 | 3 | - | - | - | - | - | - | - | - | - | - | - | - | - | - | - |
|  | *Huea* sp. 1 | 8 | 4 | 5 | - | - | - | - | - | - | - | - | - | - | - | - | - | - |
|  | *Lecanora* cf. *mons-nivis* | 2 | - | - | - | - | - | - | 1 | - | - | - | - | - | - | - | - | - |
|  | *Lecanora fuscobrunnea* | - | 2 | - | - | - | - | - | - | - | - | - | - | - | - | 1 | - | - |
|  | *Lecanora* sp. 1 | - | - | 1 | - | - | - | - | - | - | - | - | - | - | - | - | - | - |
|  | *Lecanora* sp. 2 | 3 | 2 | - | 1 | - | - | - | - | - | - | - | - | - | - | - | - | - |
|  | *Lecanora* sp. 3 | 3 | 1 | - | - | - | 1 | - | - | - | - | - | 1 | - | - | - | - | - |
|  | *Lecidea andersonii* | - | - | 1 | - | - | - | - | - | - | - | - | - | - | - | - | - | - |
|  | *Lecidea cancriformis* | - | 1 | 2 | - | 2 | - | - | - | - | - | - | - | - | 3 | - | 7 | 1 |
|  | *Lecidea lapicida* | 1 | - | - | - | - | - | - | - | - | - | - | - | - | - | - | - | - |
|  | *Lecidea polypycnidophora* | 4 | 5 | - | - | - | - | - | - | - | - | - | - | - | - | - | - | - |
|  | *Lecidea* sp. 5 | - | - | - | - | - | - | - | - | 1 | - | - | - | - | - | - | - | - |
|  | *Lecidea* sp. 6 | 3 | - | - | - | - | - | - | - | - | - | - | - | - | - | - | - | - |
|  | *Lecidea* UCR1 | 1 | - | - | - | - | - | - | - | - | - | 1 | - | - | - | - | - | - |
|  | *Lecidella greenii* | 16 | 9 | 1 | 2 | - | - | - | - | - | - | - | - | - | - | - | 1 | - |
|  | *Rhizoplaca macleanii* | 12 | 29 | 2 | 3 | 3 | - | 1 | 1 | - | - | - | - | - | - | - | - | - |

## Online Resource 2

**Online Resource 2a**: Foehn event in Miers Valley during March 2008

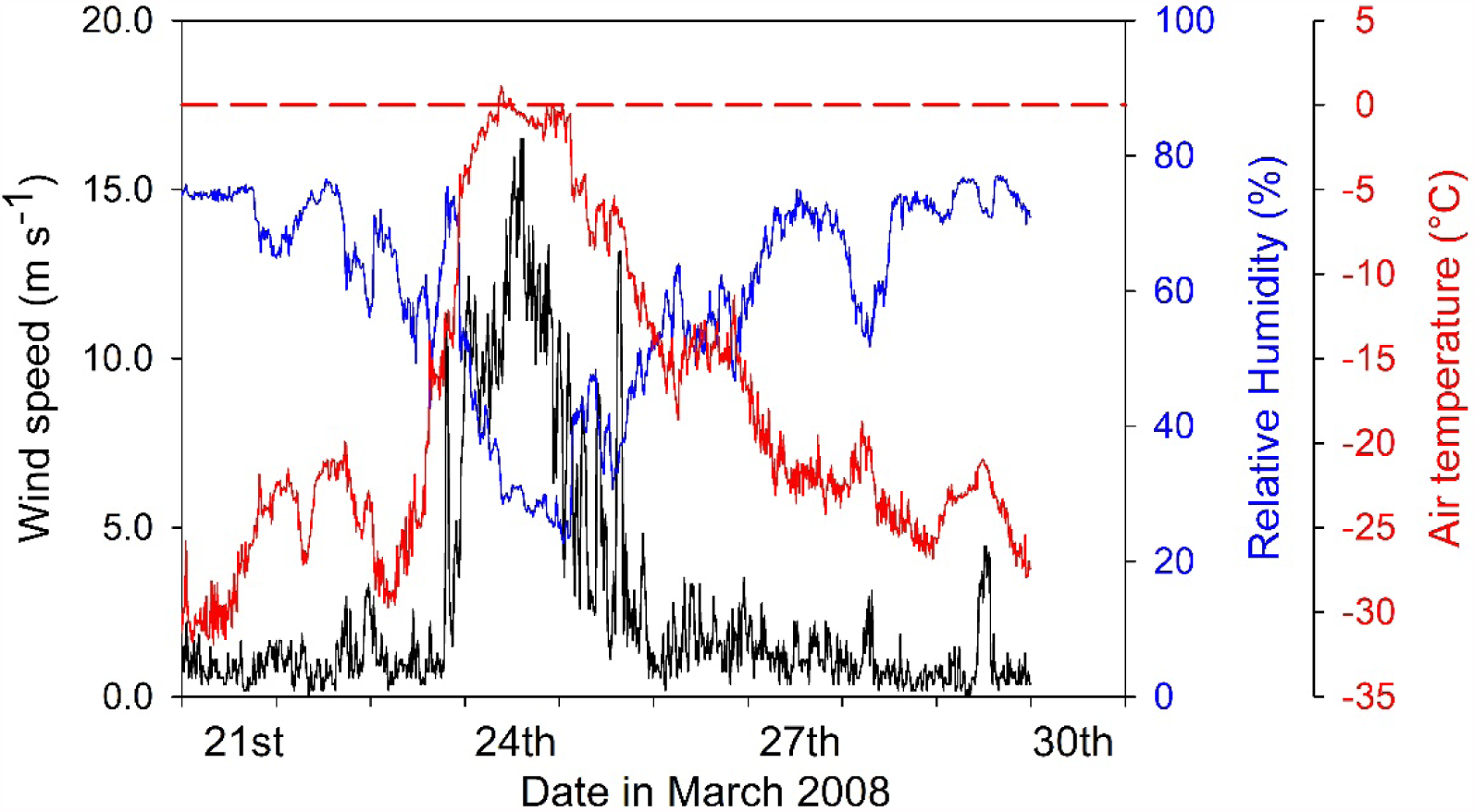

**Online Resource 2b**: Phylogeny of all mycobionts specimen based on the marker ITS and calculated with IQ-tree. Numbers in italic refer to SH-aLRT and UFboot supports. Branches with SH-aLRT < 80 % and UFboot < 95 % were collapsed.

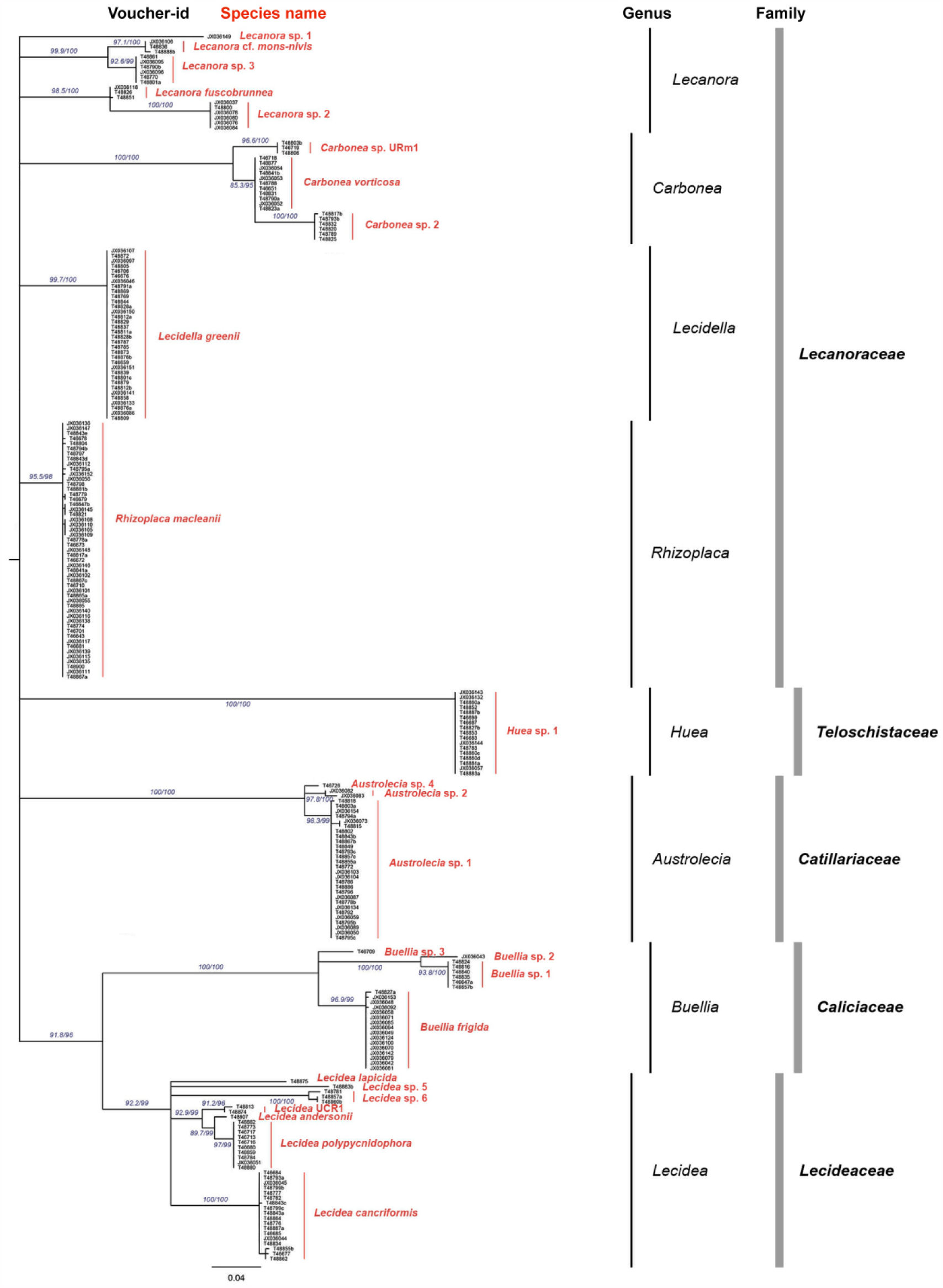

**Online Resource 2c**: Phylogeny of all photobiont specimen based on the marker ITS and calculated with IQ-tree. Numbers in italc refer to SH-aLRT and UFboot supports. Branches with SH-aLRT < 80 % and UFboot < 95 % were collapsed.

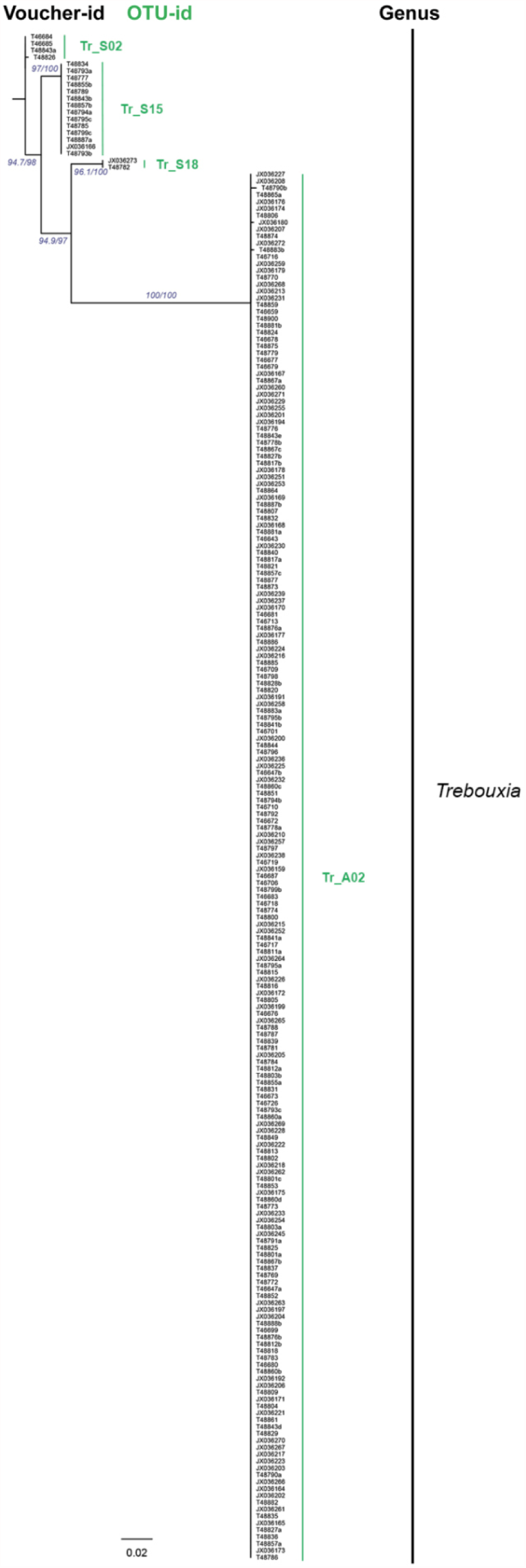

**Online Resource 2d**: Sample locations of the mycobiont species (*min10MycoSp*) and photobiont haplotypes (*min10PhoHap*) within the MDV (maps generated with http://www.gpsvisualizer.com).

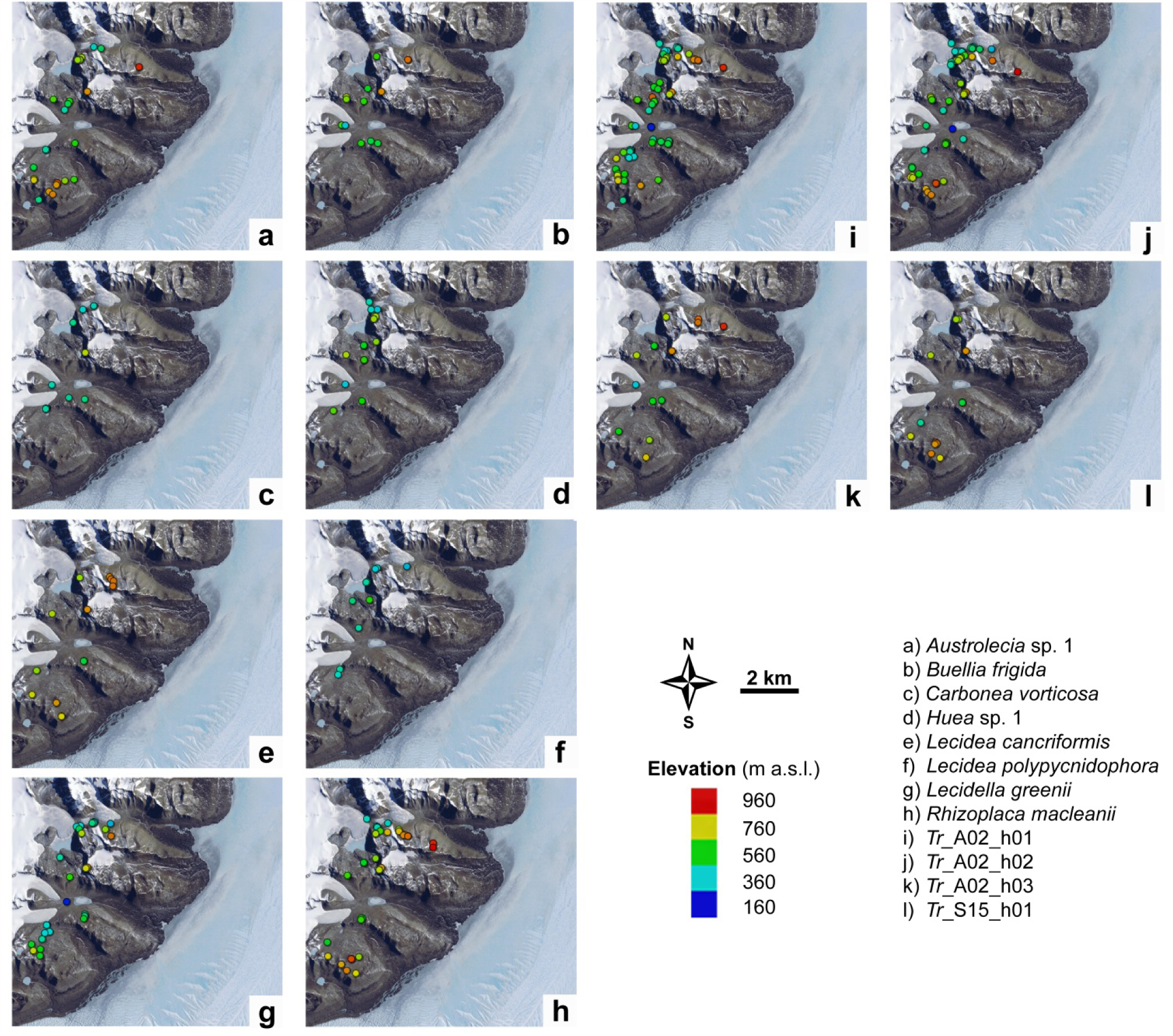

**Online Resource 2e**: Scatterplots of (a) *h / N*, (b) *Hd*, (c) *π*, (d) *d‘*, (e) *NRI*, (f) *PSV* and (g) *PSR* dependent on elevation means for *min10MycoSp. r* gives the Pearson correlation coefficent.

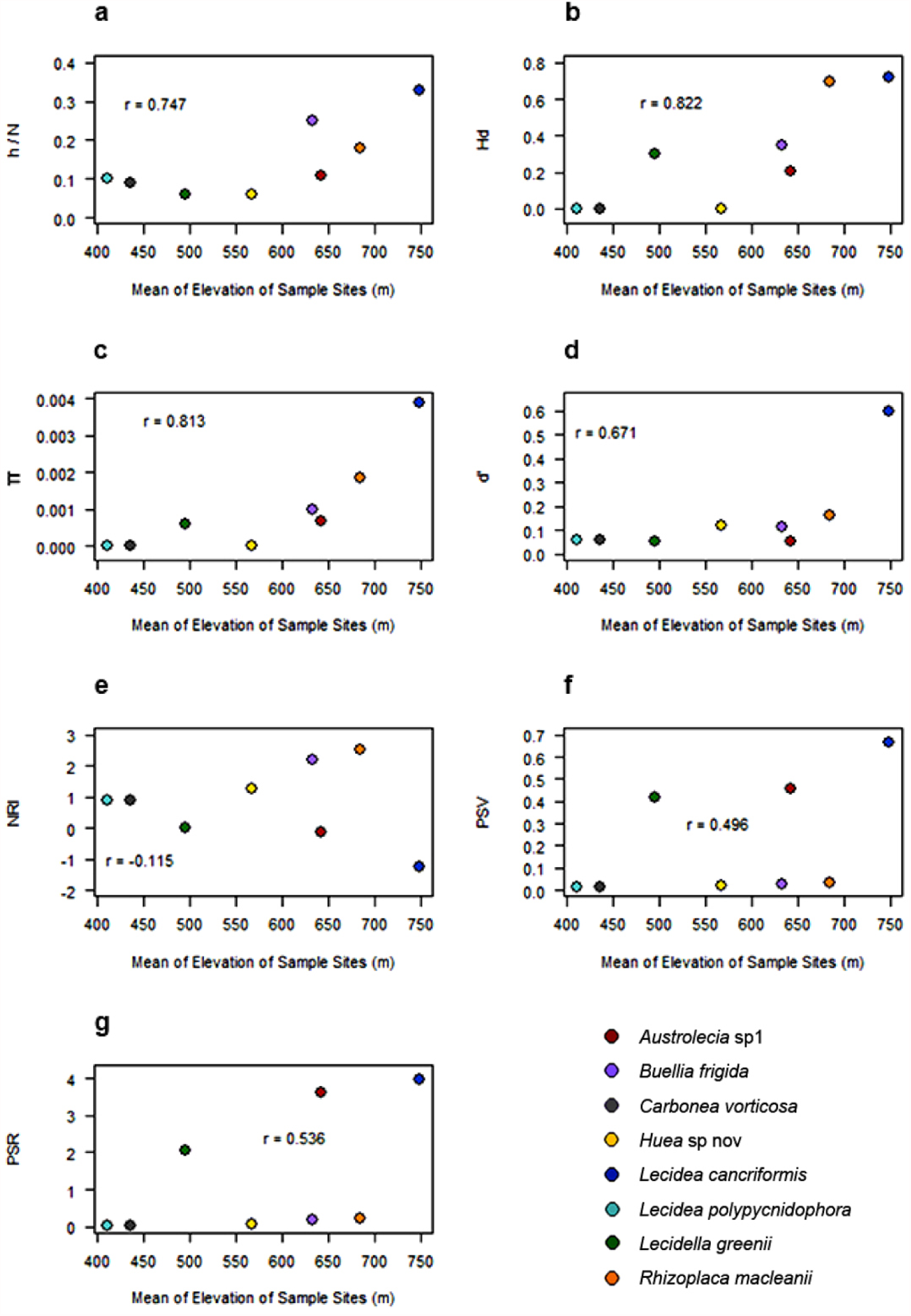

## References

Adams BJ et al. (2006) Diversity and distribution of Victoria Land biota Soil Biology & Biochemistry 38:3003–3018 doi:DOI 10.1016/j.soilbio.2006.04.030

Alatalo JM, Jagerbrand AK, Molau U (2015) Testing reliability of short-term responses to predict longer-term responses of bryophytes and lichens to environmental change Ecol Indic 58:77–85

Allen JL, Lendemer JC (2016) Climate change impacts on endemic, high-elevation lichens in a biodiversity hotspot Biodiversity and conservation 25:555–568

Ayling BF, McGowan HA (2006) Niveo-eolian sediment deposits in coastal South Victoria Land, Antarctica: Indicators of regional variability in weather and climate Arct Antarct Alp Res 38:313–324 doi:DOI 10.1657/1523-0430(2006)38[313:Nsdics]2.0.Co;2

Bassler C et al. (2016) Contrasting patterns of lichen functional diversity and species richness across an elevation gradient Ecography 39:689–698

Beck A, Kasalicky T, Rambold G (2002) Myco-photobiontal selection in a Mediterranean cryptogam community with *Fulgensia fulgida* New Phytologist 153:317–326

Blaha J, Baloch E, Grube M (2006) High photobiont diversity associated with the euryoecious lichen-forming ascomycete *Lecanora rupicola* (Lecanoraceae, Ascomycota) Biol J Linn Soc 88:283–293

Blüthgen N, Menzel F, Blüthgen N (2006) Measuring specialization in species interaction networks BMC Ecol 6:9 doi:10.1186/1472-6785-6-9

Bottos EM, Woo AC, Zawar-Reza P, Pointing SB, Cary SC (2014) Airborne Bacterial Populations Above Desert Soils of the McMurdo Dry Valleys, Antarctica Microb Ecol 67:120–128

Castello M (2003) Lichens of Terra Nova Bay area, northern Victoria Land (Continental Antarctica) Studia Geobotanica:3–54

Colesie C, Gommeaux M, Green TGA, Budel B (2014a) Biological soil crusts in continental Antarctica: Garwood Valley, southern Victoria Land, and Diamond Hill, Darwin Mountains region Antarct Sci 26:115–123

Colesie C, Green TGA, Haferkamp I, Budel B (2014b) Habitat stress initiates changes in composition, CO2 gas exchange and C-allocation as life traits in biological soil crusts Isme J 8:2104–2115

Dal Grande F, Rolshausen G, Divakar PK, Crespo A, Otte J, Schleuning M, Schmitt I (2018) Environment and host identity structure communities of green algal symbionts in lichens New Phytologist 217:277–289

Dal Grande F et al. (2017) Adaptive differentiation coincides with local bioclimatic conditions along an elevational cline in populations of a lichen-forming fungus Bmc Evol Biol 17 doi:ARTN 9310.1186/s12862-017-0929-8

De Los Rios A, Wierzchos J, Ascaso C (2014) The lithic microbial ecosystems of Antarctica’s McMurdo Dry Valleys Antarct Sci 26:459–477

Doran PT, McKay CP, Clow GD, Dana GL, Fountain AG, Nylen T, Lyons WB (2002) Valley floor climate observations from the McMurdo dry valleys, Antarctica, 1986–2000 J Geophys Res-Atmos 107 doi:Artn 477210.1029/2001jd002045

Dormann CF, Gruber B, Fruend J (2008) Introducing the bipartite Package: Analysing Ecological Networks R news 8/2:8–11

Ellis AR, Burchett WW, Harrar SW, Bathke AC (2017) Nonparametric Inference for Multivariate Data: The R Package npmv J Stat Softw 76:1–18 doi:10.18637/jss.v076.i04

Fernandez-Mendoza F, Domaschke S, Garcia MA, Jordan P, Martin MP, Printzen C (2011) Population structure of mycobionts and photobionts of the widespread lichen Cetraria aculeata Mol Ecol 20:1208–1232 doi:10.1111/j.1365-294X.2010.04993.x

Fountain AG, Nylen TH, Monaghan A, Basagic HJ, Bromwich D (2010) Snow in the McMurdo Dry Valleys, Antarctica Int J Climatol 30:633–642

Gardes M, Bruns TD (1993) Its Primers with Enhanced Specificity for Basidiomycetes - Application to the Identification of Mycorrhizae and Rusts Mol Ecol 2:113–118

Green AT, Schlensog M, Sancho LG, Winkler BJ, Broom FD, Schroeter B (2002) The photobiont determines the pattern of photosynthetic activity within a single lichen thallus containing cyanobacterial and green algal sectors (photosymbiodeme) Oecologia 130:191–198 doi:10.1007/s004420100800

Green TGA (2009) Lichens in arctic, antarctic and alpine ecosystems. Rundgespräche der Kommission für Ökologie. In: Ökologische Rolle der Flechten. vol 36. Verlag Dr. Friedrich Pfeil, pp 45–65

Green TGA, Sancho LG, Pintado A, Schroeter B (2011a) Functional and spatial pressures on terrestrial vegetation in Antarctica forced by global warming Polar Biol 34:1643–1656 doi:DOI 10.1007/s00300-011-1058-2

Green TGA, Sancho LG, Turk R, Seppelt RD, Hogg ID (2011b) High diversity of lichens at 84 degrees S, Queen Maud Mountains, suggests preglacial survival of species in the Ross Sea region, Antarctica Polar Biol 34:1211–1220 doi:DOI 10.1007/s00300-011-0982-5

Green TGA, Schroeter B, Sancho L (2007) Plant life in Antarctica. In: Pugnaire FI, Valladares F (eds) Functional Plant Ecology. 2nd edn. CRC Press, Boca Raton, Florida, pp 389–433

Grytnes JA, Heegaard E, Ihlen PG (2006) Species richness of vascular plants, bryophytes, and lichens along an altitudinal gradient in western Norway Acta Oecol 29:241–246

Guindon S, Dufayard JF, Lefort V, Anisimova M, Hordijk W, Gascuel O (2010) New Algorithms and Methods to Estimate Maximum-Likelihood Phylogenies: Assessing the Performance of PhyML 3.0 Syst Biol 59:307–321 doi:10.1093/sysbio/syq010

Head JW, Marchant DR (2014) The climate history of early Mars: insights from the Antarctic McMurdo Dry Valleys hydrologic system Antarct Sci 26:774–800 doi:10.1017/S0954102014000686

Helmus MR, Bland TJ, Williams CK, Ives AR (2007) Phylogenetic measures of biodiversity Am Nat 169:E68–E83

Henskens FL, Green TGA, Wilkins A (2012) Cyanolichens can have both cyanobacteria and green algae in a common layer as major contributors to photosynthesis Ann Bot-London 110:555–563 doi:DOI 10.1093/Aob/Mcs108

Hertel H (1984) Über saxicole, lecideoide Flechten der Subantarktis Beiheft zur Nova Hedwigia 79:399–499

Hertel H (1987) Progress and problems in taxonomy of Antarctic saxicolous lecideoid lichens Bibliotheca Lichenologica 25:219–242

Hertel H (1998) Flechten im Hochgebirge. In: Jung WW (ed) Naturerlebnis Alpen (Jubiläumsschrift zum 50-jährigen Bestehen der naturkundlichen Abteilung der Sektion München im Deutschen Alpenverein E.V. Verlag Dr. F. Pfeil, pp 33–48

Hertel H (2007) Notes on and records of Southern Hemisphere lecideoid lichens Bibliotheca Lichenologica 95:267–296

Hertel H (2009) A new key to cryptothalline species of the genus Lecidea (Lecanorales) Bibliotheca Lichenologica 99:185–204

Junker RR, Larue-Kontic AAC (2018) Elevation predicts the functional composition of alpine plant communities based on vegetative traits, but not based on floral traits Alpine Bot 128:13–22 doi:10.1007/s00035-017-0198-6

Kalyaanamoorthy S, Minh BQ, Wong TKF, von Haeseler A, Jermiin LS (2017) ModelFinder: fast model selection for accurate phylogenetic estimates Nat Methods 14:587-+

Kappen L (2000) Some aspects of the great success of lichens in Antarctica Antarct Sci 12:314–324

Kappen L, Valladares F (2007) Opportunistic growth and desiccation tolerance: the ecological success of poikilohydrous autotrophs Functional Plant Biology

Katoh K, Misawa K, Kuma K, Miyata T (2002) MAFFT: a novel method for rapid multiple sequence alignment based on fast Fourier transform Nucleic Acids Res 30:3059–3066 doi:DOI 10.1093/nar/gkf436

Kearse M et al. (2012) Geneious Basic: An integrated and extendable desktop software platform for the organization and analysis of sequence data Bioinformatics 28:1647–1649 doi:10.1093/bioinformatics/bts199

Keller I, Alexander JM, Holderegger R, Edwards PJ (2013) Widespread phenotypic and genetic divergence along altitudinal gradients in animals J Evolution Biol 26:2527–2543 doi:10.1111/jeb.12255

Kembel SW et al. (2010) Picante: R tools for integrating phylogenies and ecology Bioinformatics 26:1463–1464

Konietschke F, Placzek M, Schaarschmidt F, Hothorn LA (2015) nparcomp: An R Software Package for Nonparametric Multiple Comparisons and Simultaneous Confidence Intervals Journal of Statistical Software 64

Körner C (2003) Alpine plant life: functional plant ecology of high mountain ecosystems. 2nd edn. Springer, Berlin, London

Körner C (2007) The use of ‘altitude’ in ecological research Trends Ecol Evol 22:569–574 doi:10.1016/j.tree.2007.09.006

Kroken S, Taylor JW (2000) Phylogenetic species, reproductive mode, and specificity of the green alga Trebouxia forming lichens with the fungal genus Letharia Bryologist 103:645–660

Lange OL (1997) Flechten in Wüsten, Halbwüsten und Steppen. In: Schöller H (ed) Geschichte, Biologie, Systematik, Ökologie, Naturschutz und kulturelle Bedeutung, vol Kleine Senckenberg-Reihe 27. Senckenbergische Naturforschende Gesellschaft, Frankfurt am Main, pp pp. 129–138

Lange OL (2000) Die Lebensbedingungen von Bodenkrusten-Organismen: Tagesverlauf der Photosynthese einheimischer Erdflechten*) Hoppea, Denkschr Regensb Bot Ges 61:423–443

Lange OL, Kilian E, Ziegler H (1986) Water-Vapor Uptake and Photosynthesis of Lichens - Performance Differences in Species with Green and Blue-Green-Algae as Phycobionts Oecologia 71:104–110

Leavitt SD et al. (2015) Fungal specificity and selectivity for algae play a major role in determining lichen partnerships across diverse ecogeographic regions in the lichen-forming family Parmeliaceae (Ascomycota) Mol Ecol 24:3779–3797 doi:10.1111/mec.13271

Leavitt SD, Nelsen MP, Lumbsch HT, Johnson LA, St Clair LL (2013) Symbiont flexibility in subalpine rock shield lichen communities in the Southwestern USA Bryologist 116:149–161 doi:10.1639/0007-2745-116.2.149

Lee CK et al. (2019) Biotic interactions are an unexpected yet critical control on the complexity of an abiotically driven polar ecosystem Commun Biol 2 doi:ARTN 6210.1038/s42003-018-0274-5

Levy J (2013) How big are the McMurdo Dry Valleys? Estimating ice-free area using Landsat image data Antarct Sci 25:119–120 doi:10.1017/S0954102012000727

Librado P, Rozas J (2009) DnaSP v5: a software for comprehensive analysis of DNA polymorphism data Bioinformatics 25:1451–1452 doi:10.1093/bioinformatics/btp187

Lindgren H, Velmala S, Hognabba F, Goward T, Holien H, Myllys L (2014) High fungal selectivity for algal symbionts in the genus Bryoria Lichenologist 46:681–695

Matheny PB, Liu YJJ, Ammirati JF, Hall BD (2002) Using RPB1 sequences to improve phylogenetic inference among mushrooms (Inocybe, Agaricales) American journal of botany 89:688–698

McKay CP (2015) Testing the Doran summer climate rules in Upper Wright Valley, Antarctica Antarct Sci 27:411–415 doi:10.1017/S095410201500005x

Mckendry IG, Lewthwaite EWD (1990) The Vertical Structure of Summertime Local Winds in the Wright Valley, Antarctica Bound-Lay Meteorol 51:321–342

Minh BQ, Nguyen MAT, von Haeseler A (2013) Ultrafast Approximation for Phylogenetic Bootstrap Mol Biol Evol 30:1188–1195 doi:10.1093/molbev/mst024

Monaghan AJ, Bromwich DH, Powers JG, Manning KW (2005) The climate of the McMurdo, Antarctica, region as represented by one year of forecasts from the Antarctic Mesoscale Prediction System J Climate 18:1174–1189

Muggia L, Perez-Ortega S, Kopun T, Zellnig G, Grube M (2014) Photobiont selectivity leads to ecological tolerance and evolutionary divergence in a polymorphic complex of lichenized fungi Ann Bot-London 114:463–475 doi:10.1093/aob/mcu146

Nadyeina O, Dymytrova L, Naumovych A, Postoyalkin S, Werth S, Cheenacharoen S, Scheidegger C (2014) Microclimatic differentiation of gene pools in the Lobaria pulmonaria symbiosis in a primeval forest landscape Mol Ecol 23:5164–5178

Nei M (1987) Molecular Evolutionary Genetics. Columbia University Press, New York

Nei M, Li WH (1979) Mathematical-Model for Studying Genetic-Variation in Terms of Restriction Endonucleases P Natl Acad Sci USA 76:5269–5273 doi:10.1073/pnas.76.10.5269

Nelsen MP, Gargas A (2009) Symbiont flexibility in Thamnolia vermicularis (Pertusariales: Icmadophilaceae) Bryologist 112:404–417

Ochyra R, R.I. LS, H. B-O (2008) The illustrated moss flora of Antarctica. Cambridge University Press, Cambridge

Otalora MAG, Martinez I, O’Brien H, Molina MC, Aragon G, Lutzoni F (2010) Multiple origins of high reciprocal symbiotic specificity at an intercontinental spatial scale among gelatinous lichens (Collemataceae, Lecanoromycetes) Molecular phylogenetics and evolution 56:1089–1095 doi:10.1016/j.ympev.2010.05.013

Pannewitz S et al. (2005) Photosynthetic responses of three common mosses from continental Antarctica Antarct Sci 17:341–352 doi:DOI 10.1017/S0954102005002774

Paradis E (2010) pegas: an R package for population genetics with an integrated-modular approach Bioinformatics 26:419–420 doi:10.1093/bioinformatics/btp696

Peksa O, Skaloud P (2011) Do photobionts influence the ecology of lichens? A case study of environmental preferences in symbiotic green alga Asterochloris (Trebouxiophyceae) Mol Ecol 20:3936–3948 doi:DOI 10.1111/j.1365-294X.2011.05168.x

Perez-Ortega S, Ortiz-Alvarez R, Allan Green TG, de Los Rios A (2012) Lichen myco- and photobiont diversity and their relationships at the edge of life (McMurdo Dry Valleys, Antarctica) FEMS Microbiol Ecol 82:429–448 doi:10.1111/j.1574-6941.2012.01422.x

Pérez-Ortega S, Ortiz-Álvarez R, Green TGA, de los Ríos A (2012) Lichen myco- and photobiont diversity and their relationships at the edge of life (McMurdo Dry Valleys, Antarctica) FEMS Microbiol Ecology 82:429–448 doi:10.1111/j.1574-6941.2012.01422.x

Pointing SB, Chan YK, Lacap DC, Lau MCY, Jurgens JA, Farrell RL (2009) Highly specialized microbial diversity in hyper-arid polar desert P Natl Acad Sci USA 106:19964–19969 doi:10.1073/pnas.0908274106

Puillandre N, Lambert A, Brouillet S, Achaz G (2012) ABGD, Automatic Barcode Gap Discovery for primary species delimitation Mol Ecol 21:1864–1877

Romeike J, Friedl T, Helms G, Ott S (2002) Genetic diversity of algal and fungal partners in four species of Umbilicaria (Lichenized ascomycetes) along a transect of the Antarctic peninsula Mol Biol Evol 19:1209–1217

Ruprecht U, Brunauer G, Printzen C (2012a) Genetic diversity of photobionts in Antarctic lecideoid lichens from an ecological viewpoint Lichenologist 44:661–678 doi:10.1017/S0024282912000291

Ruprecht U, Brunauer G, Türk R (2014) High photobiont diversity in the common European soil crust lichen Psora decipiens Biodiversity and conservation 23:1771–1785 doi:10.1007/s10531-014-0662-1

Ruprecht U, Fernandez-Mendoza F, Türk R, Fryday A (2019) High levels of endemism and local differentiation in the fungal and algal symbionts of saxicolous lecideoid lichens along a latitudinal gradient in southern South America BioRxiv doi: 10.1101/699942

Ruprecht U, Lumbsch HT, Brunauer G, Green TGA, Turk R (2012b) Insights into the Diversity of Lecanoraceae (Lecanorales, Ascomycota) in continental Antarctica (Ross Sea region) Nova Hedwigia 94:287–306 doi:10.1127/0029-5035/2012/0017

Ruprecht U, Lumbsch HT, Brunauer G, Green TGA, Türk R (2010) Diversity of Lecidea (Lecideaceae, Ascomycota) species revealed by molecular data and morphological characters Antarct Sci 22:727–741 doi:10.1017/S0954102010000477

Sancho LG, Pintado A, Green TGA (2019) Antarctic Studies Show Lichens to be Excellent Biomonitors of Climate Change Diversity 11:42

Sancho LG et al. (2017) Recent Warming and Cooling in the Antarctic Peninsula Region has Rapid and Large Effects on Lichen Vegetation Sci Rep-Uk 7

Schlensog M, Schroeter B, Sancho LG, Pintado A, Kappen L (1997) Effect of strong irradiance on photosynthetic performance of the melt-water dependent cyanobacterial lichen Leptogium puberulum (Collemataceae) Hue from the maritime Antarctic. In: Kappen L (ed) Bibliotheca Lichenologica: New Species and Novel Aspects in Ecology and Physiology of Lichens. Berlin, pp 235–246

Schroeter B, Green TGA, Pannewitz S, Schlensog M, Sancho LG (2010) Fourteen degrees of latitude and a continent apart: comparison of lichen activity over two years at continental and maritime Antarctic sites Antarct Sci 22:681–690

Schroeter B, Green TGA, Pannewitz S, Schlensog M, Sancho LG (2011) Summer variability, winter dormancy: lichen activity over 3 years at Botany Bay, 77 degrees S latitude, continental Antarctica Polar Biol 34:13–22

Seppelt RD, Türk R, Green TGA, Moser G, Pannewitz S, Sancho LG, Schroeter B (2010) Lichen and moss communities of Botany Bay, Granite Harbour, Ross Sea, Antarctica Antarct Sci 22:691–702 doi:DOI 10.1017/S0954102010000568

Singh G, Dal Grande F, Divakar PK, Otte J, Crespo A, Schmitt I (2017) Fungal-algal association patterns in lichen symbiosis linked to macroclimate New Phytologist 214:317–329

Speirs JC, Steinhoff DF, McGowan HA, Bromwich DH, Monaghan AJ (2010) Foehn Winds in the McMurdo Dry Valleys, Antarctica: The Origin of Extreme Warming Events Journal of Climate 23:3577–3598 doi:10.1175/2010jcli3382.1

Stichbury G, Brabyn L, Green TGA, Cary C (2011) Spatial modelling of wetness for the Antarctic Dry Valleys Polar Res 30 doi:Artn 6330 DOI 10.3402/Polar.V30i0.6330

Trifinopoulos J, Nguyen LT, von Haeseler A, Minh BQ (2016) W-IQ-TREE: a fast online phylogenetic tool for maximum likelihood analysis Nucleic Acids Res 44:W232–W235

Vargas Castillo R, Beck A (2012) Photobiont selectivity and specificity in Caloplaca species in a fog-induces community in the Atacama Desert, northern Chile Fungal Biology 116:665–676

Wagner M, Trutschnig W, Bathke AC, Ruprecht U (2017) A first approach to calculate BIOCLIM variables and climate zones for Antarctica Theoretical and Applied Climatology:1–19 doi:10.1007/s00704-017-2053-5

Webb CO (2000) Exploring the phylogenetic structure of ecological communities: An example for rain forest trees Am Nat 156:145–155 doi:DOI 10.1086/303378

Webb CO, Ackerly DD, McPeek MA, Donoghue MJ (2002) Phylogenies and community ecology Annu Rev Ecol Syst 33:475–505 doi:10.1146/annurev.ecolysis.33.010802.150448

Werth S, Sork VL (2010) Identity and genetic structure of the photobiont of the epiphytic lichen Ramalina menziesii on three oak species in Southern California American journal of botany 97:821–830 doi:DOI 10.3732/Ajb.0900276

White TJ, Bruns TD, Lee SB, Taylor JW (1990) Amplification and direct sequencing of fungal ribosomal RNA Genes for phylogenies. In: Innis MA, Gelfand DH, Sninsky JJ, White TJ (eds) PCR Protocols: A Guide to Methods and Applications. Academic Press, San Diego, pp 315–322

Wirtz N, Lumbsch HT, Green TGA, Turk R, Pintado A, Sancho L, Schroeter B (2003) Lichen fungi have low cyanobiont selectivity in maritime Antarctica New Phytologist 160:177–183 doi:DOI 10.1046/j.1469-8137.2003.00859.x

Wolf JHD (1993) Diversity Patterns and Biomass of Epiphytic Bryophytes and Lichens Along an Altitudinal Gradient in the Northern Andes Ann Mo Bot Gard 80:928–960

Wornik S, Grube M (2010) Joint dispersal does not imply maintenance of partnerships in lichen symbioses Microb Ecol 59:150–157 doi:DOI 10.1007/s00248-009-9584-y

Yahr R, Vilgalys R, DePriest PT (2006) Geographic variation in algal partners of Cladonia subtenuis (Cladoniaceae) highlights the dynamic nature of a lichen symbiosis. (vol 171, pg 847, 2006) New Phytologist 172:377–377

Yung CCM et al. (2014) Characterization of Chasmoendolithic Community in Miers Valley, McMurdo Dry Valleys, Antarctica Microb Ecol 68:351–359 doi:10.1007/s00248-014-0412-7

Zahlbruckner A (1925) Catalogus Lichenum Universalis. In, vol III. Leipzig,

Zoller S, Scheidegger C, Sperisen C (1999) PCR primers for the amplification of mitochondrial small subunit ribosomal DNA of lichen-forming ascomycetes Lichenologist 31:511–516

